# Molecular determinants of Hrp1–RNA recognition underlying yeast RNA Polymerase II transcription attenuation

**DOI:** 10.64898/2026.06.16.732720

**Authors:** Catalina Lujan-Rodriguez, Max A. Popoloski, Lane E. Couturier, Jessica J. Richa, Justin M. Talluto, Madison E. Lapine, Mackenzie Roche, Sidney J. Edouard, Vincent Pavan, Jason N. Kuehner

## Abstract

Premature termination of transcription (PTT), also known as attenuation, is a conserved gene regulatory mechanism that operates across all domains of life and in viruses. Attenuation enables rapid cellular responses to environmental and metabolic changes and fine-tunes expression of biosynthetic genes. In *Saccharomyces cerevisiae*, attenuation of RNA Polymerase II (Pol II) transcription was first linked to the Nrd1-Nab3-Sen1 (NNS) termination pathway for non-coding RNAs, and the mRNA 3’-end processing factor Hrp1 has been implicated more recently. Substitutions in Hrp1 RNA Recognition Motifs (RRMs) cause attenuator readthrough and reduce RNA-binding affinity *in vitro*, but direct evidence for Hrp1 functioning at attenuators *in vivo* remains limited. Here, we characterized 5’-end RNA terminator elements from several genes, including *RAD3*, *SNG1*, *MNR2*, and *CPR8*. Readthrough mutations clustered in AU-rich regions resembling polyadenylation site (pA) efficiency elements, consistent with Hrp1 binding targets. Amino acid substitutions of Hrp1 RRM residue F162 revealed a general requirement for aromaticity in RNA recognition that varied to some degree by gene context. To test Hrp1-RNA interactions independent of other yeast factors, we adapted a bacterial 3-hybrid (B3H) assay. Hrp1 interacted with RNA derived from the *GAL7* 3’-end pA site and 5’-end terminator regions of *RAD3*, *MNR2*, and *CPR8*. Mutations in AU-rich RNA regions that disrupted Pol II attenuation in yeast generally impaired B3H interactions. However, some Hrp1 mutants (M191T, I270T, D271G, M275V, T280I) retained binding to *CPR8* terminator RNA, suggesting their defects require additional yeast components. These results demonstrate that Hrp1 is sufficient to bind multiple UA-rich attenuator RNAs *in vivo*, expanding Hrp1 function to include early transcription events.

## INTRODUCTION

RNA Polymerase II (Pol II) synthesizes all protein-coding genes in eukaryotes and is subject to regulation at all three stages of transcription, including initiation, elongation, and termination (Archuleta et al. 2024; Kuldell and Kaplan 2025). During transcription termination, the recognition of RNA elements by protein factors contributes to the release of Pol II and its nascent transcript from the DNA template. Pol II termination downstream of protein-coding genes is coupled with 3’-end mRNA processing; thus, a polyadenylation (pA) site serves a critical signaling role (Kaur et al. 2021; Rodríguez-Molina et al. 2023; Martinez and Svejstrup 2025). Premature Pol II termination (PTT) at upstream regions of protein-coding genes (i.e. attenuation) can also occur, with possible roles for regulation and/or RNA quality control (Kamieniarz-Gdula and Proudfoot 2019; Rouvière et al. 2022; Bentley 2024). Instead of producing full-length mRNA, Pol II transcription attenuation can result in a truncated RNA that encodes an incomplete protein, lacks a coding sequence, and/or is rapidly degraded.

Transcription attenuation has been observed in all domains of life and viruses, demonstrating its biological utility. In bacteria, where the mechanism of PTT was first identified, attenuation enables rapid cellular responses to environmental and metabolic changes and fine-tunes expression of biosynthetic genes (Turnbough 2019). In archaea, promoter-proximal recruitment of termination factor aCPSF1 contributes to attenuation, which can be modulated in response to oxidative damage (Blombach et al. 2021; Pilotto and Werner 2022). In eukaryotes, attenuators are linked to developmental regulation and control of stress response genes (Cugusi et al. 2022; Menon et al. 2024; Lysakovskaia et al. 2025; Tayari and Shiekhattar 2026). Attenuation represses HIV-1 viral gene expression in latently infected cells, which supports immune evasion and impedes antiretroviral therapy (Said et al. 2023). Furthermore, attenuator readthrough has been linked to several human diseases (Grzechnik and Mischo 2025).

Pol II attenuation has been best studied in the model eukaryote yeast *Saccharomyces cerevisiae*. Several examples of attenuator-based autoregulation have been described, including the *NRD1, HRP1,* and *PCF11* genes, which encode Pol II termination and 3’-end RNA processing factors (Arigo et al. 2006; Steinmetz et al. 2006; Kuehner and Brow 2008; Creamer et al. 2011; Grzechnik et al. 2015). Other yeast targets of Pol II attenuation include genes involved in nucleotide biosynthesis (*IMD2*, *URA2*), cell wall damage signaling (*FKS2*), glucose starvation (*CLN3*), glycogen metabolism (*GPH1*), and nitrogen metabolism (*GLT1*) (Jenks et al. 2008; Kuehner and Brow 2008; Kwapisz et al. 2008; Thiebaut et al. 2008; Kim and Levin 2011; Darby et al. 2012; Chen et al. 2017; Merran and Corden 2017). In these cases, the primary pathway responsible for attenuator recognition involves the RNA-binding proteins Nrd1 and Nab3 and the RNA/DNA helicase Sen1. The Nrd1/Nab3/Sen1 (NNS) pathway also functions in transcription termination at many non-coding RNAs (e.g. snRNAs, snoRNAs, cryptic unstable transcripts) (Arndt and Reines 2015; Chaves-Arquero and Pérez-Cañadillas 2023). The integrator complex in higher eukaryotes bears similarity to the yeast NNS pathway given its involvement in snRNA 3’-end processing and promoter-proximal termination (Wagner et al. 2023; Tayari and Shiekhattar 2026).

In contrast to NNS-dependent attenuators, our lab has identified targets that rely on the mRNA 3’-end cleavage factor (CF), cleavage and polyadenylation factor (CPF), and Sen1, without appreciable Nrd1/Nab3 involvement (Graber et al. 2013; Whalen et al. 2018). The DNA repair factor gene *DEF1* exhibits a 5’-end “hybrid” attenuation mechanism (Whalen et al. 2018), which is reminiscent of the 3’-end crosstalk of CPF-CF and NNS complexes during failsafe termination (Lemay and Bachand 2015). In cases of attenuator recognition where Nrd1/Nab3 involvement is limited, we hypothesized that cleavage factor IB (CFIB/Hrp1) serves a critical RNA-binding role (Amodeo et al. 2022). *DEF1* attenuator readthrough mutations clustered in a *DEF1* 5’-end region matching a yeast pA site efficiency element (consensus: 5’-UAUAUA-3’) (Amodeo et al. 2022), a preferred Hrp1 binding site (Kessler et al. 1997; Valentini et al. 1999; Kaur et al. 2021). In addition, substitutions in the Hrp1 RNA recognition motif (RRM) disrupted *DEF1* attenuator recognition (Amodeo et al. 2022). Additional Hrp1-dependent targets in yeast genes *RAD3*, *SNG1*, and *MNR2* were identified based on several attenuator hallmarks, including: (1) upstream (5’-end) peaks of Pol II occupancy with reduced signal throughout the open reading frame (Schaughency et al. 2014), (2) promoter-proximal pA sites (Johnson et al. 2011), (3) upstream enrichment of Hrp1 (Tuck and Tollervey 2013) and (4) upstream depletion of Nrd1 and Nab3 (Jamonnak et al. 2011). For all three candidates, conditional degradation of Hrp1 resulted in attenuator readthrough (Amodeo et al. 2022).

Our previous research on Hrp1-dependent Pol II attenuation suggested several areas for additional study. As a 3’-end processing factor, Hrp1 is known to mediate both protein-RNA and protein-protein interactions (Barnwal et al. 2012), but its required function(s) for 5’-end attenuation are less clear. *DEF1* attenuation requires UA-rich RNA elements, and it is unknown if this is a general requirement for Hrp1-dependent PTT. The Hrp1 RRMs are highly conserved across RNA-binding proteins in yeast and other eukaryotes, including the human Hrp1 ortholog HNRNPDL (Goguen and Brow 2023). Mutations in HNRNPDL are associated with limb-girdle muscular dystrophy type 3 (Vieira et al. 2014). Substitutions in RRM1 residue Hrp1-F162 disrupt the function of both *DEF1* and *HRP1* 5’-end mRNA attenuators (Amodeo et al. 2022; Goguen and Brow 2023), and other RRM mutants decrease Hrp1 affinity for (UA)_4_ RNA *in vitro* (Goguen et al. 2026). The Hrp1-F162 residue also supports 3’-end Pol II termination of the *SNR82* non-coding RNA (Goguen and Brow 2023; Goguen et al. 2026), raising the question of what is common and distinct among 5’-and 3’-end terminators.

In this study, we employed a yeast genetic selection to identify mutations that promote Pol II readthrough across several Hrp1-dependent attenuators, which frequently targeted conserved UA-rich RNA elements. We investigated the biochemical determinants of Hrp1 RRM interaction with attenuator RNA, revealing a preference for aromatic residues at position F162. In addition, we optimized a bacterial B3H system to assay Hrp1-RNA interactions in the absence of other yeast factors, revealing gene-specific differences based on terminator RNA sequence and/or protein context.

## METHODS

### Gibson cloning of terminator sequences into reporter plasmids

For yeast reporter gene assays, terminator regions were amplified using Q5 Hot Start High-Fidelity 2X Master Mix (New England Biolabs) from yeast genomic DNA template and cloned into pGAC24-noT-*CUP1* and/or pGAC24-noT-*lacZ* plasmids (*LEU2*) using the HiFi DNA Assembly Cloning Kit (New England Biolabs). Briefly, primers were designed to amplify the following upstream regions relative to the start codon: *CPR8* (−107 to +164), *DEF1* (−187 to +93), *HRP1* (−419 to +80), *MNR2* (−225 to +116), *RAD3* (−139 to +175), and *SNG1* (−262 to +115). For downstream regions, the following sequences were amplified relative to the stop codon: *CYC1* (+113 to +205), and *GAL7* (+15 to +252). For the non-coding *SNR82* gene, the downstream region +269 to +468 was amplified relative to the transcription start site. Primers were designed using NEBuilder to contain appropriate homology to the pGAC24-noT-*CUP1* and/or pGAC24-noT-*lacZ* plasmids, which were digested with restriction enzyme XhoI and phosphatase-treated with Antarctic phosphatase (New England Biolabs) prior to assembly. Following transformation into 5-alpha competent cells (New England Biolabs), candidates were screened via colony PCR using GoTaq DNA Polymerase (Promega), and candidates containing inserts were verified by Sanger sequencing (QuintaraBio). Transformation of reporter plasmids into yeast strains was performed using a standard lithium acetate method.

The pRS313-(3x)HA-(6x)Gly-HRP1 (*HIS3*) construct was generated using the HiFi DNA Assembly Cloning Kit (New England Biolabs). A gBlock DNA fragment containing 3x(HA)-6x(Gly) was synthesized, containing homology to the upstream region of the *HRP1* ORF (Integrated DNA Technologies). Primers were designed using NEBuilder to amplify the pRS313-*HRP1* (HIS3) plasmid (Whalen et al. 2018) in two fragments, with appropriate homology to the gBlock fragment. Following transformation into 5-alpha competent cells (New England Biolabs), candidates were verified by Sanger sequencing (QuintaraBio).

For bacterial reporter gene assays, terminator regions or *HRP1* regions were amplified from plasmid DNA templates (see above) or yeast genomic DNA template and cloned into pBait (pSS2-1xMS2^hp^-7GC-T_trpA_; AddGene 222406) (Nguyen et al. 2025) using SmaI/HindIII restriction sites or pPrey (pBRα-β flap 831-105; AddGene 53734) using NotI/BamHI restriction sites and the HiFi DNA Assembly Cloning Kit (New England Biolabs). Primers were designed using NEBuilder to contain appropriate homology to target plasmids.

Following restriction digestion and prior to assembly, plasmids were treated with Antarctic phosphatase (New England Biolabs). Following transformation into 5-alpha competent cells for pBait plasmids or 5-alpha F’I^q^ competent cells for pPrey plasmids (New England Biolabs), plasmids were purified with a ZR plasmid miniprep kit (Zymo) and candidates were verified by Sanger sequencing (QuintaraBio).

### Mutagenesis of terminator RNAs and Hrp1 proteins

Random mutagenesis of the 5’-end terminator regions in *CPR8*, *MNR2*, *RAD3*, and *SNG1* was carried out by error-prone PCR and *in-vivo* homologous recombination (Muhlrad et al. 1992). Primers complementary to the relevant pGAC24-terminator-*CUP1* plasmid sequences were used to direct amplification by *Taq* DNA polymerase (NEB) of a fragment containing the terminator region along with vector homology. PCR amplification was performed for 40 rounds, purified, and repeated for an additional 40 rounds, relying upon the normal error rate of *Taq* DNA polymerase for the introduction of random mutations.

The pGAC24-terminator-*CUP1* plasmids were digested with *Bam*HI and *Xho*I, creating a gap flanked by complementarity to the PCR product. Digested plasmids were treated with Antarctic phosphatase (NEB), purified using a Zymoclean Gel DNA Recovery kit (Zymo Research), and cotransformed with the PCR product (5:1 insert:vector ratio) into the 46α yeast strain. After selection for transformants on SC-Leu plates, colonies were replica plated onto SC-Leu plates containing copper sulfate (0.2 or 0.4 mM CuSO_4_). Plasmids were recovered from copper-resistant colonies and retransformed into yeast to confirm that the copper-resistant phenotype was linked to the plasmid prior to Sanger sequencing (QuintaraBio).

Site-directed mutagenesis of terminator regions in bacterial B3H pBait plasmids, *HRP1* in yeast plasmid pRS313-(3x)HA-(6x)Gly-*HRP1*, or bacterial B3H plasmid pPrey-*HRP1* (1-534) was performed using primers designed with QuikChange Primer Design software and the Quikchange Lightning site-directed mutagenesis kit (Agilent) or NEBaseChanger software and the Q5 site-directed mutagenesis kit (NEB). Following transformation into 5-alpha competent cells for pBait or pRS313 plasmids and 5-alpha F’I^q^ competent cells for pPrey plasmids (New England Biolabs), candidates were verified by Sanger sequencing (QuintaraBio).

### Bacterial B3H lacZ assays

B3H assays were performed as described in (Stockert et al. 2022), with some alterations indicated below. Competent cells of the FW102-OL_2_-62 bacterial strain were made with the Mix and Go transformation kit (Zymo Research) and one of the three B3H plasmids (e.g. pAdapter) pre-transformed so that only two plasmids needed to be transformed in at the same time (e.g. pPrey + pBait). Empty vectors of pBait, pPrey, or pAdapter were included as negative controls where needed. Competent cells (100 uL aliquots) were freshly transformed with 150 ng of each B3H plasmid of interest (5 uL of 30 ng/uL stock) and selected on LB plates supplemented with kanamycin (50 ug/mL), carbenicillin (100 ug/mL), chloramphenicol (25 ug/mL), and spectinomycin (100 ug/mL), grown overnight at 37°C. Single colonies (n=3-4) were selected from transformation plates and grown in 200 uL of B3H liquid media, which was LB liquid medium supplemented with carbenicillin (100 ug/mL), chloramphenicol (25 ug/mL), kanamycin (50 ug/mL), spectinomycin (100 ug/mL), and arabinose (0.2% wt/vol) in a 96-well plate. The plate was sealed with breathable film (VWR), and cultures were grown in a shaking incubator at 700 RPM and 37°C or 30°C overnight. The following day, saturated overnight cultures were diluted in fresh B3H media (20:200) and grown in a 96-well plate with a plastic lid at 700 RPM and 37°C for 1.5 to 2 hours or 30°C for 2.5 to 3 hours until mid-log phase OD_600_ = 0.2 to 0.4. Absorbance was measured using a SpectraMax 190 microplate reader (Molecular Devices), and OD_600_ readings of individual wells were kept as close to each other as possible (ideally within 0.2 OD_600_ units). Lysis was performed by adding 20 uL of PopCulture reagent (Millipore) and 0.05 uL of T4 lysozyme (NEBExpress) per reaction (prepared as a mastermix) and incubating at 700 RPM and 37°C for 20 minutes. During the incubation, a fresh Z-buffer mixture was prepared by adding 120 uL of Z-buffer (60 mM Na_2_HPO_4_*7H_2_O, 40 mM NaH_2_PO_4_*H_2_O, 10 mM KCl, 1 mM MgSO_4_, pH 7.0), 30 uL of ONPG (4 mg/mL), and 0.3 uL of β-mercaptoethanol per reaction (prepared as a mastermix). The Z-buffer mixture (150 uL) was added into a new section of the 96-well plate, followed by 30 uL of lysed cells.

Absorbance was measured at OD_420_ using a SpectraMax 190 microplate reader (Molecular Devices), with the temperature set to 28°C and readings taken every minute for one hour. The β-Gal activity was calculated in Miller units according to the formula:

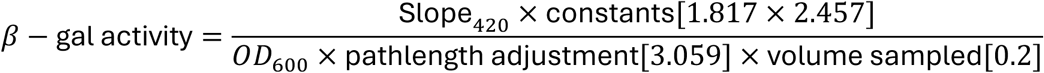

Fold-stimulation was calculated by dividing the mutant activity by the wild-type activity.

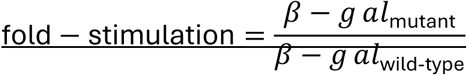

Statistical testing was performed using Graphpad Prism version 11 for MacOS and Welch’s ANOVA (\**P* ≤ 0.05, \*\**P* ≤ 0.01, \*\*\**P* ≤ 0.001, \*\*\*\**P* ≤ 0.0001, ns—not significant).

### Bacterial B3H spot test assays

For bacterial spot test assays, mid-log phase cells (OD_600_ = 0.2 to 0.4) were diluted 2:198 with B3H liquid media in a 96-well plate. All samples were mixed and spotted using a 96-well replica plater (Sigma) on LB plates supplemented with kanamycin (50 ug/mL), carbenicillin (100 ug/mL), chloramphenicol (25 ug/mL), spectinomycin (100 ug/mL), arabinose (0.2% wt/vol), X-Gal (40 ug/mL), phenylethyl-β-D-thiogalactopyranoside (400 uM), and IPTG (1.5 uM). The plates were air-dried and incubated overnight at 37°C, followed by 4°C overnight storage and imaging with a digital camera and DigiBox (Emmanuel College).

### Yeast terminator-lacZ assays

Single colonies (n=3) were selected from transformation plates and grown overnight in 5 mL of appropriate selective liquid media (e.g. SC -His/-Leu) in test tubes in a shaking incubator at 30°C. Cultures were diluted 20:180 and absorbance at OD_600_ was measured for each strain using a SpectraMax 190 microplate reader (Molecular Devices). Cultures were diluted to OD_600_ = 0.15 to 0.30 in a 200 uL volume in a 96-well plate. Cell lysis was performed and reporter enzyme activity was measured using the Yeast β-Galactosidase Assay Kit according to manufacturer’s instructions (Thermo Scientific). The assay kit working solution was prepared and added separately to the 96-well plate in 100 µL aliquots. The absorbance at OD_420_ was measured every minute for one hour to collect a kinetic reaction rate and slope. Slopes were gathered using a kinetic reaction window in a linear range with a strong R^2^ value. Relative β-galactosidase activity was calculated as previously described (Thibodeau *et al*. 2004).

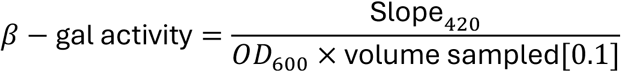

### Yeast spot test assays

Yeast bearing the reporter plasmids of interest were grown overnight in a test tube shaking incubator at 30°C in selective liquid media (US Biological). Absorbance at OD_600_ was measured using a spectrophotometer (Implen Diluphotometer) and cultures were diluted to OD_600_ = 1.0. Each diluted culture (200 μL) was transferred to a 96-well plate, followed by 10-fold serial dilutions (x4) into adjacent wells. All samples were mixed and spotted using a 96-well replica plater (Sigma). For *CUP1* strains, spots were plated on SC -Leu plates or SC -Leu plates + copper sulfate (0.2 mM) (US Biological). For *HA-HRP1*/*AID-HRP1* strains, spots were plated on SC-His plates −/+ synthetic auxin (500 uM α-Naphthaleneacetic Acid; NAA). The plates were air-dried and incubated for 2-4 days, followed by imaging with a digital camera and DigiBox (Emmanuel College).

### Western blot analysis of proteins from yeast and bacterial extracts

To detect HA-Hrp1 proteins, the relevant pRS313-*HA-HRP1(HIS3)*/*AID-HRP1* yeast strains were grown overnight in a test tube shaking incubator at 30°C in SC -His selective liquid media (US Biological), followed by harvesting 5 ODs and freezing pellets at −80°C. Protein extracts were prepared using 2 M LiOAc and 0.4 M NaOH to permeabilize the yeast cell wall prior to extraction with 1× SDS-PAGE sample buffer (Zhang 2011).

To detect ⍺-NTD-Hrp1 proteins in bacteria, the relevant B3H strains were grown overnight in a test tube shaking incubator at 37°C in B3H selective media, which was LB liquid medium supplemented with carbenicillin (100 ug/mL), chloramphenicol (25 ug/mL), kanamycin (50 ug/mL), spectinomycin (100 ug/mL), and arabinose (0.2% wt/vol). Absorbance at OD_600_ was measured using a spectrophotometer (Implen Diluphotometer) and 1.5 ODs of cells were harvested in safe-lock tubes and frozen at −20°C. Cells were resuspended in 100 uL of sterile water, followed by an equal volume (100 uL) of 2X SDS sample buffer (containing beta-mercaptoethanol) (Bio-Rad). Samples were heated in a 100°C heatblock for 3 minutes, followed by 4-5 brief vortexing sessions. The heat treatment and vortexing step was repeated again until solution appeared clear.

Protein extracts (10 µl) were loaded into a 10% SDS-PAGE mini-protean TGX stain-free gel (Bio-Rad), and total protein was visualized and quantified using the ChemiDoc Imaging System (Bio-Rad). Proteins were transferred to a PVDF membrane using a Trans-blot Turbo instrument (Bio-Rad), followed by blocking in 5% milk/TBST for 1 hour at room temperature. The membrane was incubated for 1 hour with either mouse anti-HA primary antibody diluted 1:1,000 in 1% milk/TBST (Santa Cruz Biotechnology, sc-7392) or mouse anti-α-NTD primary antibody (NeoClone, W0030) diluted 1:5,000 in 1% milk/TBST. The membrane was washed in 1X TBST and incubated with anti-mouse secondary antibody (diluted 1:15,000 in 1% milk/TBST; Jackson ImmunoResearch) for 1 hour at room temperature. Target proteins were visualized using Clarity chemiluminescent substrate (Bio-Rad) and a ChemiDoc Imaging System (Bio-Rad). The scanner images were uploaded into Image Lab software (Bio-Rad), and the band intensities were quantified. The signal from HA-Hrp1 or α-NTD-Hrp1 bands was divided by the signal of the total protein to normalize each lane, followed by normalization to appropriate wild-type controls.

### Molecular Modeling of RRM mutations in the context of an Hrp1-RNA structure

PyMol software (Version 3.1.6.1) was used to access an Hrp1-RNA structure (PDB: 2CJK). Using the sequence viewer setting, the structure was color-coded (Hrp1 RRM1 – green, Hrp1 RRM 2 – blue, Linker – grey, RNA – gold). The RNA was shown as a cartoon, with the settings for bases and sugars set to filled ring (with border). Amino acids corresponding to mutation sites were selected, shown as ball and stick, color-coded to purple, and main chain and hydrogen atoms were hidden.

### Multiple sequence alignment of terminator regions

Intergenic sequences were obtained from the *<u>Saccharomyces sensu stricto</u>* <u>genus</u> <u>genomes</u>, including *S. cerevisiae*, *S. paradoxus*, *S. mikatae*, *S. bayanus*, and *S. kudriavzevii* (Scannell et al. 2011). Multiple sequence alignments with hierarchical clustering were performed using <u>Multalin software</u> (Corpet 1988).

## RESULTS

### Readthrough mutations in 5’-end attenuator candidates map to conserved UA-rich RNA regions, consistent with Hrp1-binding sites

Our lab has previously identified several yeast genes (*RAD3*, *SNG1,* and *MNR2*) with hallmarks of transcription attenuation (Amodeo et al. 2022), and more recently we identified a region upstream of *CPR8* with similar characteristics. To further characterize these attenuator candidates and the RNA sequences required for their function, we utilized a yeast *CUP1* reporter gene system whereby attenuator recognition or readthrough leads to growth differences on agar media plates containing copper (**Fig. 1A**). If an attenuator is functional the *CUP1* gene is transcribed at a lower level, resulting in copper sensitivity. If a mutation leads to attenuator readthrough, the *CUP1* gene is transcribed at a higher level, resulting in copper resistance. As expected, strains containing a *CUP1* plasmid lacking a terminator (No Term, negative control) were copper resistant (**Fig. 1B-E, row 1**), and strains containing wild-type 5’-end sequences from all four candidate attenuators were copper sensitive (**Fig. 1B-E, row 2**). Error-prone PCR was used to introduce random mutations into the candidate attenuators, and a range of copper-resistant strains were identified (**Fig. 1B-E**).

**Figure 1.**
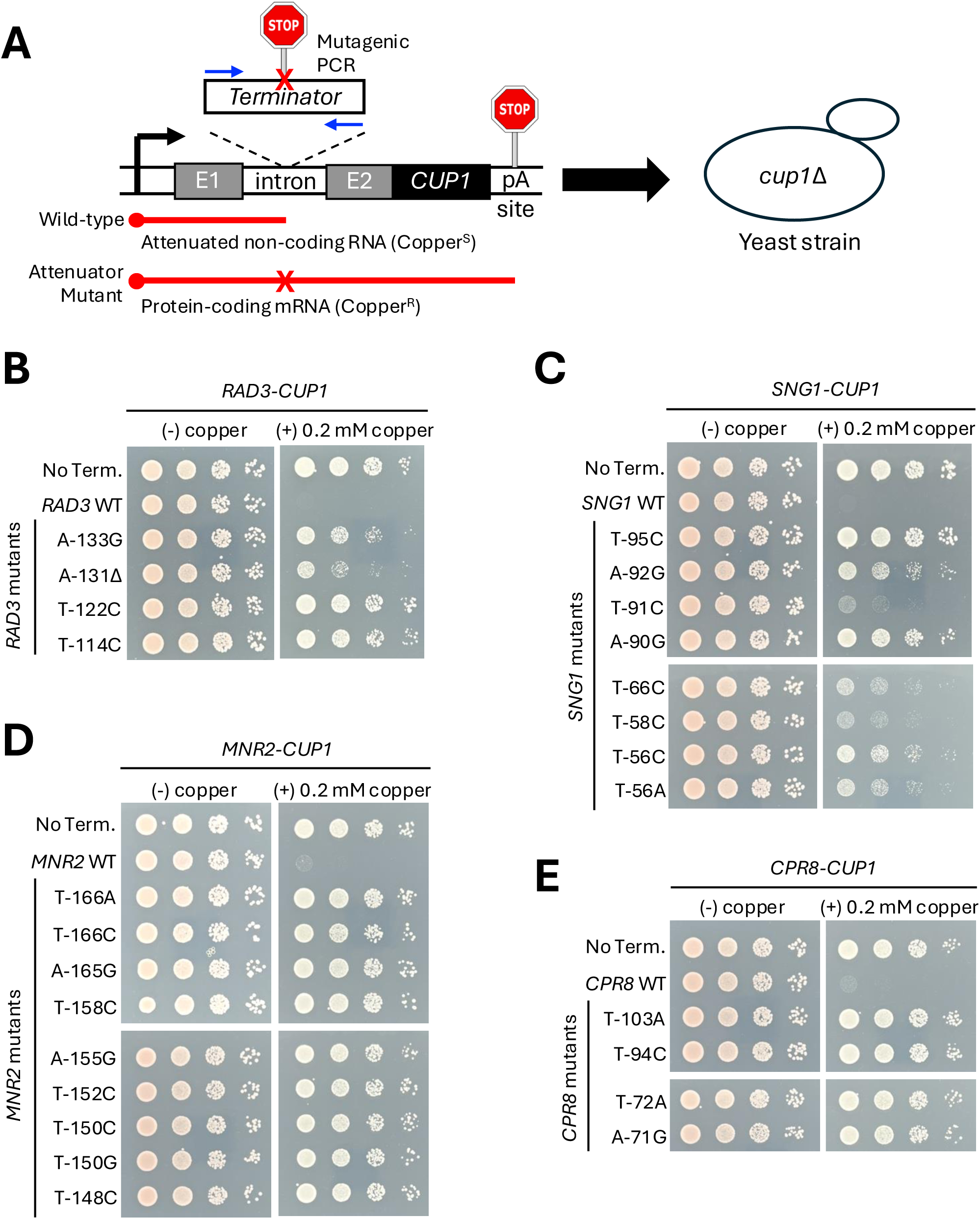
Identification of yeast Pol II transcription attenuator readthrough mutations. **(A)** Schematic of genetic selection strategy used to identify attenuator readthrough mutations. Transcription terminator regions upstream of *RAD3*, *MNR2*, *SNG1*, and *CPR8* genes were amplified via error-prone PCR and co-transformed with a gapped plasmid containing the *CUP1* reporter gene in a *cup1*Δ strain. Copper^S^ = copper sensitive, Copper^R^ = copper resistant. **(B)** Growth of yeast strains containing reporter plasmids plated on −/+ copper plates after serial dilution. Higher levels of copper-resistance indicate higher levels of attenuator readthrough. No Term. = No terminator.

To gain further insight into the sequence context of attenuator RNA function, copper-resistant point mutations were mapped to 5’-end regions of candidate genes (**Fig. 2A-D**). Sequence conservation of related yeast species was used to highlight regions that may bear functional significance. Attenuator readthrough mutations clustered in TA-rich regions (UA-rich in RNA) at least once for every candidate and were often found in multiple locations, consistent with possible roles as efficiency elements (consensus 5’-UAUAUA-3’) and Hrp1-binding sites (Kessler et al. 1997; Valentini et al. 1999; Kaur et al. 2021). Many of these TA-rich regions were conserved across different yeast species. Similar TA-rich regions have previously been identified as efficiency elements in the 3’-end pA site/terminator regions of *GAL7* and *CYC1* (**Fig. S1**) (Guo and Sherman 1995) and the 5’-end attenuator regions of *DEF1* and *HRP1* (**Fig. S2**) (Whalen et al. 2018; Goguen et al. 2026), many of which also exhibit phylogenetic conservation. For the most part, readthrough mutants were located downstream of transcription start sites (TSS) and within the expected 5’-UTR of attenuator candidates *RAD3*, *SNG1*, and *MNR2* (**Fig. 2A, 2B, 2C**). For the *CPR8* candidate, the position of readthrough mutants relative to the TSS was more complicated (**Fig. 2D**), suggesting that the more upstream region serves as a 3’-end pA site/terminator for the tandemly transcribed gene (*BUD17*) while the more downstream region may function as a 5’-end *CPR8* attenuator. Given this ambiguity, the *CPR8* region in this study will be referred to more generally as a terminator rather than an attenuator.

**Figure 2.**
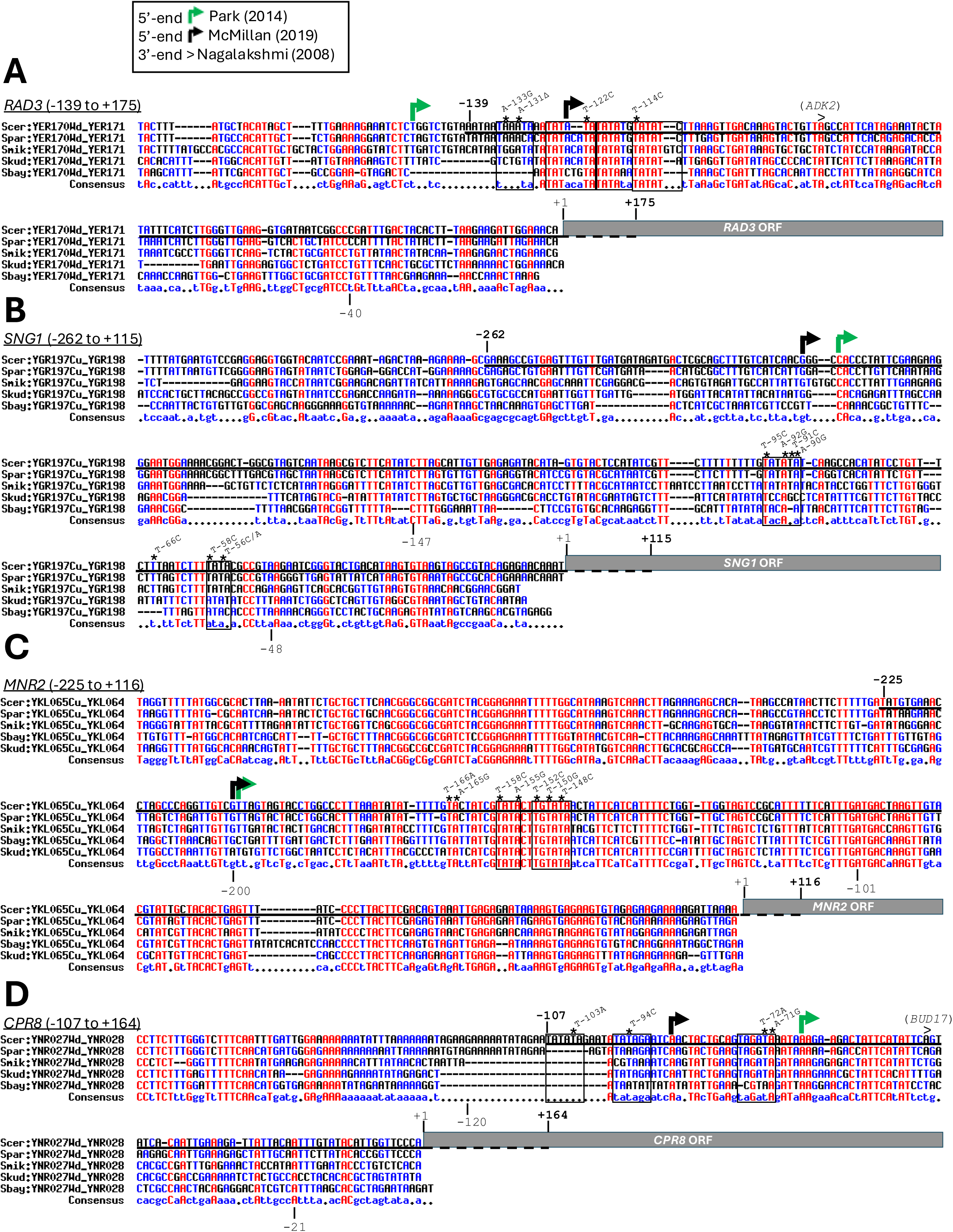
Phylogenetic analysis reveals DNA conservation in promoter-proximal Pol II transcription terminators (attenuators). DNA regions upstream of (A) *RAD3*, (B) *SNG1*, (C) *MNR2*, and (D) *CPR8* were obtained from *sensu stricto* yeast species (*S. cerevisiae*, *S. paradoxus*, *S. mikatae*, *S. bayanus*, *S. kudriavzevii*) (Scannell et al. 2011) and aligned using MultAlin software (Corpet 1988). A base that is highly conserved appears in high-consensus color (red) and as an uppercase letter in the consensus line. A base that is weakly conserved appears in a low-consensus color (blue) and as a lowercase letter in the consensus line. Other bases appears in neutral color (black). A position with no conserved base is represented by a dot in the consensus line. The 5’-end transcription start sites are indicated with arrows (black and green) (Park et al. 2014; McMillan et al. 2019), and transcription terminator regions cloned into reporter genes are underlined. The > symbol indicates the mRNA 3’-end of a neighboring gene (Nagalakshmi et al. 2008), if positioned in a tandemly transcribed orientation. Regions cloned into *CUP1* and/or *lacZ* reporter genes are underlined. Copper-resistant point mutants are indicated with asterisks and numbered relative to the respective gene +1 <u>A</u>TG start codon.

### Substitutions in Hrp1 RRM1 residue F162 reveal a preference for amino acid aromaticity in attenuator RNA attenuator recognition that varies by gene context

Given the presence of attenuator readthrough mutations in putative Hrp1-binding sites and in some cases sensitivity to Hrp1 depletion (Amodeo et al. 2022), we predicted that substitutions in the Hrp1 RNA recognition motifs (RRMs) would also be disruptive. Our lab previously identified Hrp1-F162W as a dominant negative mutant that causes Pol II readthrough for Hrp1-dependent attenuators (Amodeo et al. 2022). In the context of an Hrp1 RRM structure with an 8 nucleotide RNA (5’-A_1_U_2_A_3_U_4_A_5_U_6_A_7_U_8_’-3), the F162 side chain forms π-stacking interactions with U_5_ and A_6_ bases **(Fig. 3A**). The phenylalanine (F) amino acid has a unique combination of a hydrophobic side chain with a benzyl group, offering potential for hydrophobic (van der Waals) contacts and π-π stacking interactions with RNA. To determine if the hydrophobic and/or aromatic properties of Hrp1-F162 were more important for attenuator function, we expanded our analysis to test additional amino acid substitutions. The group one substitutions (F162A, F162I, F162L) were chosen for their exclusively hydrophobic properties and lack of aromaticity. The group two substitutions (F162W, F162H, F162Y) were chosen for their range of hydrophobicity (phenylalanine > tryptophan > tyrosine > histidine) and their aromaticity, with F, W, H, and Y side chains containing 1 ring, 2 rings, 1 ring, and 1 ring, respectively.

**Figure 3.**
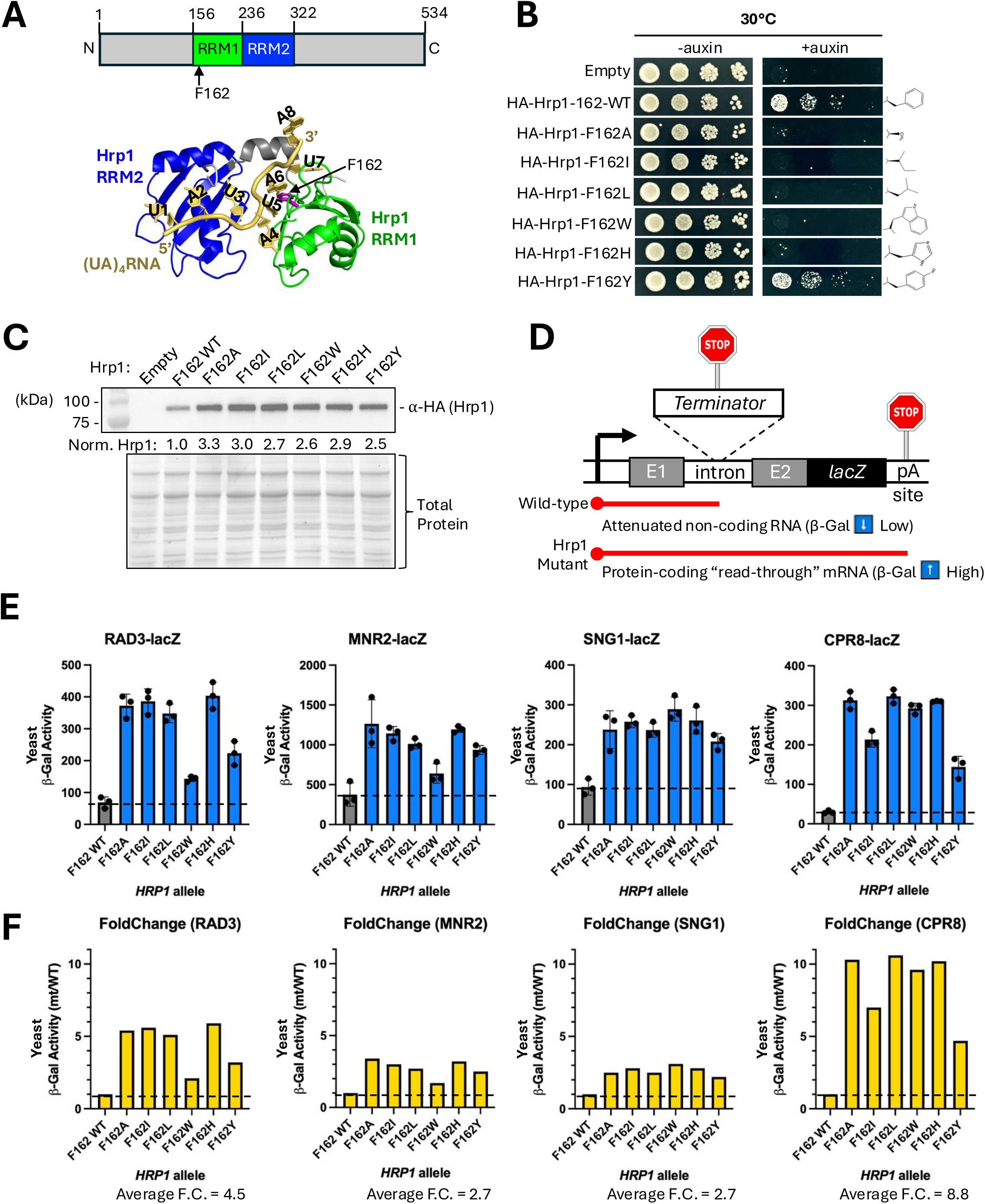
Amino acid substitutions in Hrp1 RNA Recognition Motif (RRM) residue F162 support a functional preference for aromaticity and gene-specific RNA terminator interactions. **(A)** Schematic of Hrp1 protein primary structure and its RNA recognition motifs (RRMs), with amino acid numbering indicated. The Hrp1-RNA structure (PDB: 2CJK) (Perez-Canadillas 2006) displays stacking interactions between the Hrp1-F162 RRM1 amino acid and U5/A6 RNA nucleotides. **(B)** Growth of yeast strains containing a chromosomal auxin-inducible degron (AID-Hrp1) transformed with WT or mutant HA-Hrp1 plasmids plated on −/+ synthetic auxin plates (α-Naphthaleneacetic Acid; NAA) after serial dilution. R-group chemical structures of HA-Hrp1-F162 WT and mutant amino acids are indicated at right. **(C)** Western blot analysis of HA-Hrp1 proteins expressed in AID-Hrp1 strains. Norm. Hrp1 = Normalized HA-Hrp1 protein relative to total protein relative to wild-type. **(D)** Schematic of reporter gene strategy to identify Pol II attenuator readthrough in HA-Hrp1-F162 protein mutants. Promoter-proximal transcription terminator regions from *RAD3*, *MNR2*, *SNG1*, and *CPR8* genes were cloned upstream of a *lacZ* reporter gene. Attenuator recognition = Low β-galactosidase expression, Attenuator readthrough = High β-galactosidase expression. **(E)** Yeast strains bearing HA-Hrp1 WT and mutant plasmids and terminator-*lacZ* reporter gene plasmids were grown to saturation overnight at 30°C and diluted prior to cell lysis and incubation with ONPG substrate, using absorption at OD_600_ for cell density and OD_420_ for β-gal production. Experiments were performed in biological triplicate, and error bars show standard deviation. **(F)** Fold-change was calculated by dividing mutant β-gal values by their respective wild-type β-gal values.

To test the viability of Hrp1-F162 mutants in yeast, a chromosomal auxin-inducible degron (AID-Hrp1) strain (Amodeo et al. 2022) was transformed with HA-Hrp1-WT and HA-Hrp1-mutant plasmids and grown on −/+ auxin plates (**Fig. 3B**). As expected, the AID-Hrp1 strain transformed with an empty vector plasmid was inviable +auxin due to the depletion of AID-Hrp1 (**Fig. 3B, row 1**), and the AID-Hrp1 strain transformed with the Hrp1-F162 wild-type (WT) plasmid was viable +auxin (**Fig. 3B, row 2**). The Hrp1-F162A, I, L, W, and H mutant strains were inviable +auxin (**Fig. 3B, rows 3-7**), confirming their toxicity as the sole *hrp1* allele. The Hrp1-F162Y mutant strain grew similarly to Hrp1-F162-WT +auxin (**Fig. 3B, row 8**). We confirmed that all six Hrp1-F162 mutant proteins were expressed (**Fig. 3C**) so the lack of yeast viability was due to an altered Hrp1 function rather than simply absence of the protein. All Hrp1-F162 mutants were expressed at higher levels (2.5 to 3.3-fold) than Hrp1-F162-WT, consistent with Pol II readthrough of the autoregulatory *HRP1* attenuator and increased Hrp1 mRNA transcription (Goguen and Brow 2023). The class 1 mutants (A, I, L) averaged a bit more Hrp1 protein expression than class 2 mutants (W, H, Y). The Hrp1-F162Y mutant exhibited the least amount of Hrp1 over-expression (2.5-fold), suggesting slightly better *HRP1* attenuator function of F162Y versus other Hrp1 mutants. Given that tyrosine belongs to the group two class of mutants, we conclude that aromaticity (single benzyl ring) is an important functional feature of F162, with some tolerance for increased hydrophilicity.

To test the importance of Hrp1-F162 more specifically for our RNA attenuators of interest, we utilized a *lacZ* reporter gene system whereby attenuator recognition or readthrough leads to lower or higher β-galactosidase activity, respectively (**Fig. 3D**). As expected based on the yeast viability testing and *HRP1* expression assays above, group one mutants (A, I, L) were more disruptive than group two mutants (W, H, Y) for *RAD3*-lacZ and *MNR2*-lacZ reporters, averaging ∼30% more readthrough. This was due to the fact that both Hrp1-F162W and Hrp1-F162Y exhibited less β-gal activity (i.e. better attenuator function) compared to the A, I, L, and H mutants (**Fig. 3E**). The *CPR8*-lacZ reporter likewise showed a reduced impact for Hrp1-F162Y, but Hrp1-F162W was as disruptive as other mutants for this terminator. The *SNG1*-lacZ reporter was similarly affected by all Hrp1-F162 mutants, with no clear difference in readthrough defects for group one or group two substitutions. As far as overall sensitivity to Hrp1-F162 mutants, the *MNR2* and *SNG1* attenuators were least sensitive, exhibiting an average fold-change (mutant/wild-type) of 2.7 for β-gal activity (**Fig. 3F**). The *RAD3* attenuator was moderately sensitive, with an average fold-change of 4.5 (**Fig. 3F)**. The *CPR8* terminator was most sensitive, with an average fold-change of 8.8 (**Fig. 3F)**. Overall, we observed that each upstream gene terminator was responsive to F162 mutants, indicating an Hrp1-dependency, but allele-specific differences and overall sensitivity suggest that attenuator context (e.g. RNA sequence and/or protein interactors) influences recognition.

### Hrp1 is sufficient for interacting with RNAs from 3’-end pA sites and 5’-end attenuators in the absence of other yeast proteins

To determine if attenuator RNA and Hrp1 mutations were impacting their interaction directly, we leveraged a bacterial 3-hybrid (B3H) assay (Stockert et al. 2022; Nguyen et al. 2025). In this assay, attenuator RNA “bait” and Hrp1 “prey” were tested for interactions that promote lacZ reporter gene activation by bridging an adapter transcription activator complex, increasing RNA polymerase recruitment (**Fig. 4A**). The attenuator RNA “bait” was fused to the MS2 RNA hairpin, and the Hrp1 protein “prey” was fused to the α-NTD subunit of RNA Polymerase. For the pPrey-α-NTD-Hrp1 fusion protein, we created plasmids with both Hrp1 RRM alone (amino acids 156-321) and Hrp1 full-length (amino acids 1-534), and their expression in the bacterial B3H strain was confirmed by Western blot (**Fig. 4B**). For the pBait-RNA fusions, we tested terminator RNAs from the downstream *GAL7* 3’-end pA site, attenuator RNAs from the *RAD3* and *MNR2* 5’-end regions, and a terminator RNA from the *CPR8* 5’-end region. When all three B3H components (adapter, prey, and bait) were present (+APB), the β-gal activity was significantly more than negative controls, where individual components were omitted (-A, -P, -B) **(Fig. 4C)**. These quantitative results using lysed cells, ONPG substrate, and a microplate spectrophotometer were further validated using a qualitative assay with agar plates +X-gal substrate, where more blue coloration was observed in +APB cells versus negative control strains (**Fig. 4C**). B3H tests of Hrp1 prey interaction with RNA bait from other 5’-end attenuators (*DEF1* and *SNG1*) and the 3’-end pA site from *CYC1* were not significantly different from the -adapter negative control (**Fig. S3**). Varying the RNA bait size did not significantly change B3H β-gal activity (**Fig. 4D**). Varying the prey protein size and bacterial growth temperature generally increased B3H interaction when using full-length Hrp1 versus the RRM alone and 37°C versus 30°C (**Fig. 4E**, **4F**). Given the results of this B3H optimization, we used minimal RNA constructs (100 bases) with full-length Hrp1 protein in bacterial strains grown at 37°C for future assays.

**Figure 4.**
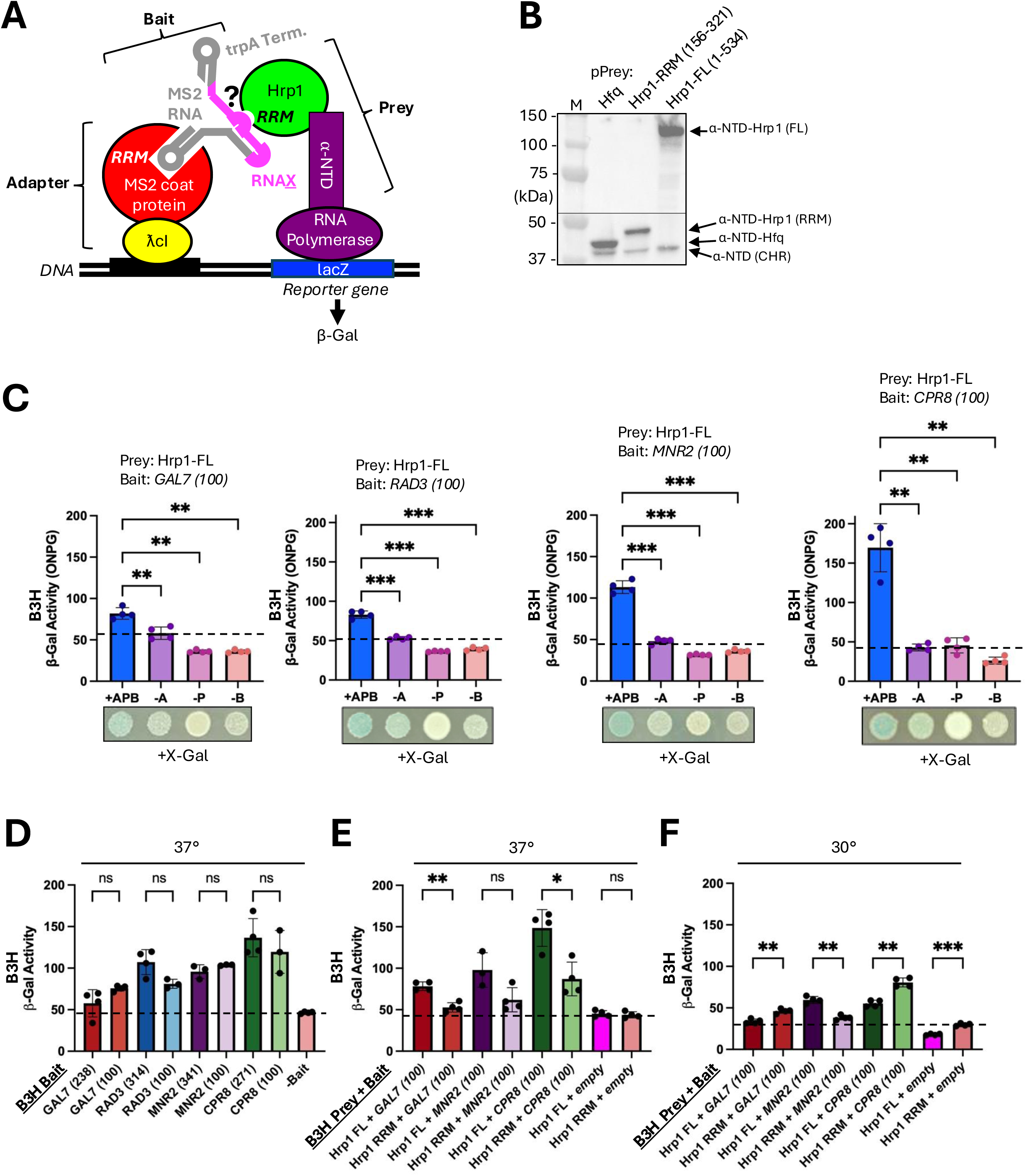
Bacterial 3-hybrid (B3H) assay detects Hrp1 protein interaction with multiple RNA transcription terminator targets. **(A)** Schematic of bacterial 3-hybrid (B3H) system, consisting of adapter, bait, and prey components (Stockert et al. 2022). The “adapter” consists of the λcI DNA-binding protein fused to the MS2 RNA-binding coat protein. The “bait” consists of the MS2 RNA hairpin fused to the RNA of interest (RNA<u>X</u>), followed by a *trpA* terminator hairpin (Nguyen et al. 2025). The “prey” consists of the Hrp1 RNA-binding protein fused to the ⍺-NTD protein, a subunit of bacterial RNA polymerase (RNAP). If Hrp1 interacts with RNA<u>X</u>, all three B3H components are bridged, increasing RNAP recruitment and transcription of the *lacZ* reporter gene and production of β-galactosidase (β-Gal). **(B)** Western blot analysis of B3H bacterial strains transformed with pPrey-Hfq, pPrey-Hrp1-RRM (156-321), and pPrey-Hrp1-FL (1-534). Protein extracts were probed with an anti-α-NTD antibody. RRM = RNA Recognition Motif. FL = Full length. α-NTD (CHR) = endogenous chromosomal gene. **(C)** Bacterial strains bearing the indicated B3H reporter gene plasmids were grown to saturation overnight at 37°C, diluted back and grown to exponential phase, followed by cell lysis and incubation with ONPG substrate, using absorption at OD_600_ for cell density and OD_420_ for β-gal production. Experiments were performed using 3-4 biological replicates, and error bars show standard deviation. +APB = +Adapter, +Prey, +Bait. -A = -Adapter. -P = -Prey. -B = -Bait. Dashed line reflects the highest level of background signal from negative controls. Asterisks indicate statistical significance by Welch’s ANOVA (*p<0.05, **p<0.01, ***p<0.001, ns – not significant, p≥0.05). The same bacterial strains were diluted and spotted onto selective media containing X-gal substrate. **(D)** Bait RNAs matching the terminator regions used in yeast *lacZ* assays from Fig. 3 were compared to smaller 100 nt constructs in the B3H assay. -Bait is lacking RNA<u>X</u> and serves as a negative control. Prey proteins of varying sizes (Hrp1 full-length vs. Hrp1 RRM) were analyzed with several bait RNAs, with bacterial strains grown at **(E)** 37°C or **(F)** 30°C. “Empty” is the pBait plasmid lacking RNA<u>X</u> and serves as a negative control.

### Mutations in attenuator RNA “bait” and Hrp1 protein “prey” disrupt B3H interactions

We predicted that attenuator RNA and Hrp1 protein readthrough mutations identified in yeast would reduce Hrp1-RNA interactions in the B3H assay. To test this hypothesis, we used site-directed mutagenesis to introduce mutations into pBait and pPrey plasmids. For the *GAL7* pA site, we targeted a previously identified efficiency element (EE) with multiple mutations (mutx5) (**Fig. S1A**), and β-gal activity was significantly reduced to near-background levels of the -Bait negative control (**Fig. 5A, red bars**). For the *RAD3* attenuator, the T-122C mutant significantly reduced β-gal activity, but T-114C remained similar to wild-type levels, even within the context of a T-122C/T-114C double mutant (**Fig. 5A, blue bars**). For the *MNR2* attenuator, the T-158C mutant was similar to wild-type, but T-152C and T-150G mutants significantly reduced β-gal activity. The loss of interaction for double mutants T-158C/T-152C and T-152C/T-150G was similar to individual mutants (**Fig. 5A, purple bars**). For the *CPR8* terminator, β-gal activity in T-94C, T-72A, and A-71G mutants was similar to wild-type (**Fig. 5A, green bars**). In the case of *GAL7*, *RAD3*, and *MNR2* terminators, the TA-rich regions (6 bases) were duplicated in tandem, whereas in the *CPR8* terminator there was a slight separation between them (**Fig. 5B**). For all four terminator candidates, the Hrp1-F162A mutant reduced β-gal activity to the background levels of the -Prey negative control (**Fig. 5C**). This reduced Hrp1-RNA interaction was not simply due to absence of the mutant protein as Hrp1-F162A was stably expressed in B3H strains (**Fig. 5D**).

**Figure 5.**
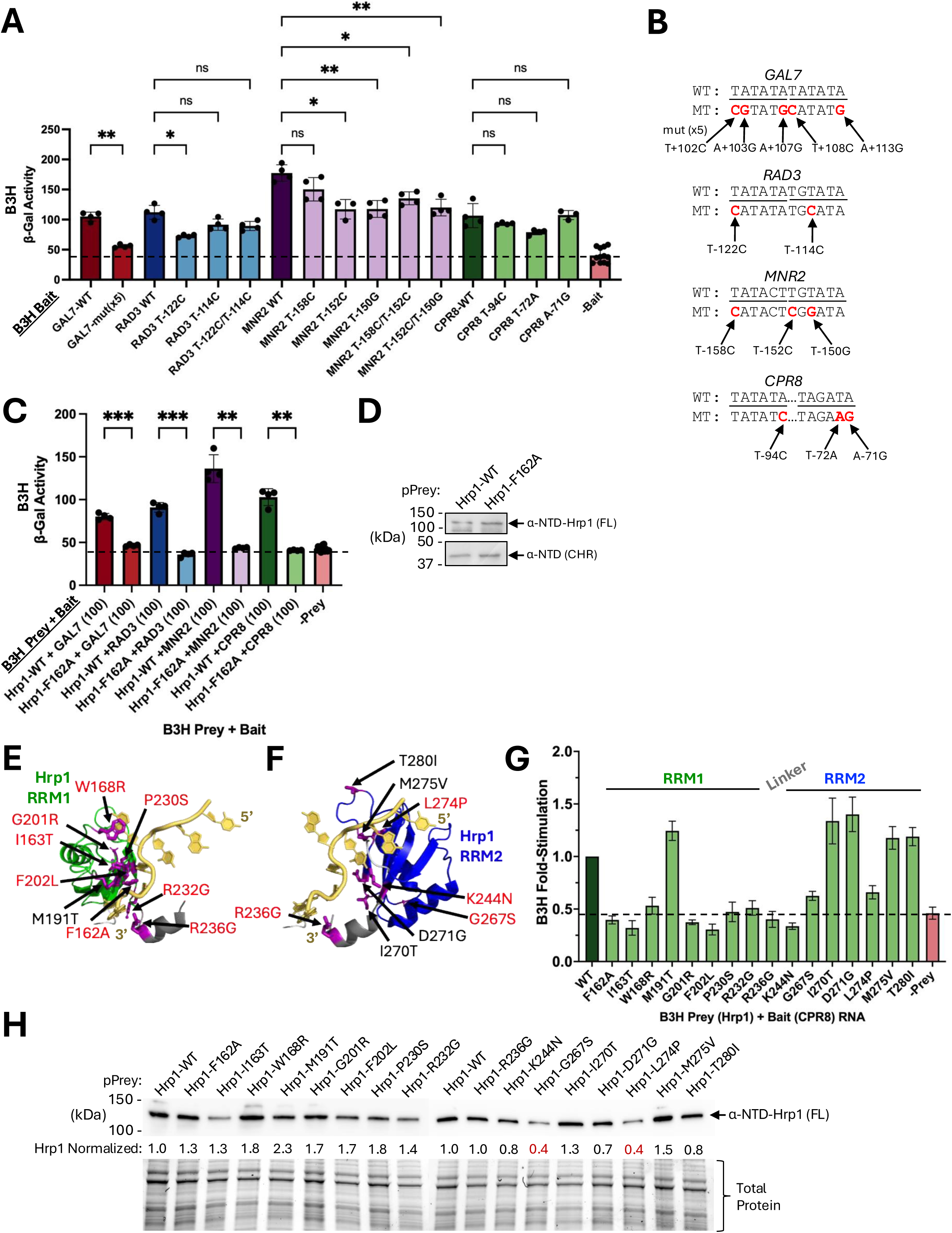
Mutations in B3H Bait and Prey components disrupt Hrp1-RNA interactions. **(A)** B3H bacterial strains were transformed with pBait-terminator constructs containing readthrough mutations identified in Figure 1, and B3H assays were performed at 37°C as described in Figure 4. -Bait is lacking RNA<u>X</u> and serves as a negative control. **(B)** pBait-terminator mutations in TA-rich DNA elements (UA-rich in RNA) are highlighted (bold/red). Underlined regions resemble the Hrp1 consensus binding site (5’-TATATA-3’). **(C)** B3H bacterial strains were transformed with pPrey-Hrp1 WT and F162A mutants, and B3H assays were performed at 37°C as described in Figure 4. -Prey is lacking Hrp1 and serves as a negative control. **(D)** Western blot analysis of B3H bacterial strains transformed with pPrey-Hrp1-WT and pPrey-Hrp1-F162A. Protein extracts were probed with an anti-α-NTD antibody. FL = Full length. α-NTD (CHR) = endogenous chromosomal gene. Hrp1-RNA structures (PDB: 2CJK) (Perez-Canadillas 2006) for **(E)** RRM1 and **(F)** RRM2 were used to visualize the locations of yeast Hrp1 readthrough mutants at the *SNR82* terminator (Goguen and Brow 2023). Hrp1 RRM mutants are highlighted to indicate if they disrupted *CPR8* terminator RNA interaction (red) or not (black). **(G)** B3H bacterial strains were transformed with pPrey-Hrp1 WT and RRM mutants, and B3H assays were performed at 37°C as described in Figure 4. -Prey is lacking Hrp1 and serves as a negative control. **(H)** Western blot analysis of B3H bacterial strains transformed with pPrey-Hrp1-WT and RRM mutants. Protein extracts were probed with an anti-α-NTD antibody. FL = Full length. Norm. Hrp1 = Normalized α-NTD-Hrp1 protein relative to total protein relative to wild-type.

### The Hrp1-dependent *SNR82* and *CPR8* terminators exhibit differential sensitivity to a subset of RRM mutants in yeast versus B3H contexts

To further dissect the importance of Hrp1 RRMs 1 and 2 in attenuator RNA recognition, we introduced a range of amino acid substitutions previously shown to cause readthrough of the *SNR82* non-coding 3’-end terminator (Goguen and Brow 2023) (**Fig. 5E**, **5F**). We tested these Hrp1 prey mutants with the *CPR8* terminator RNA bait, given the robustness of this B3H interaction compared to other candidates. As expected, most of the Hrp1 RRM mutants (F162A, I163T, W168R, G201R, F202L, P230S, R232G, R236G, K244N, G267S, L274P) reduced β-gal activity to near background levels of the -Prey negative control. Interestingly, a subset of mutants (M191T, I270T, D271G, M275V, T280I) retained β-gal activity at or above wild-type Hrp1 levels (**Fig. 5G**). For the Hrp1-G267S and Hrp1-274P mutants, their lower β-gal activity may be explained at least in part by less prey protein (levels reduced >2-fold), but the rest of the mutants were stably expressed (**Fig. 5H**). We conclude that a majority of Hrp1 readthrough mutants (11/16) for the *SNR82* non-coding 3’-end terminator likewise disrupted Hrp1-RNA interactions for the 5’-end *CPR8* terminator. For the five Hrp1 mutants that failed to disrupt the *CPR8* RNA interaction, there could be differences between *CPR8* and *SNR82* in terms of RNA sequence and/or protein context.

### The *SNR82* terminator is bipartite but more reliant on upstream sequence elements for function

To further explore the differences observed in Hrp1 mutant sensitivity for the *SNR82* non-coding 3’-end terminator and *CPR8* 5’-end terminator, we analyzed *SNR82* phylogenetic conservation and terminator behavior in yeast cells. The *SNR82* 3’-end terminator used for the identification of Hrp1 readthrough mutants includes 200 bp (**Fig. 6A**) (Goguen and Brow 2023). To further test this result in our lacZ reporter gene assay, we compared the terminator activity of *SNR82* (+269 to +468) and the upstream 100 bp (+269 to +368) versus the downstream 100 bp (+369 to +469). As expected, the 200 bp construct exhibited strong terminator activity, significantly reducing Pol II transcription by ∼12-fold compared to a *lacZ* reporter gene without an upstream terminator (No Term) (**Fig. 6B**). The *SNR82* upstream half (+269 to +369) significantly reduced Pol II termination by ∼2-fold, and the downstream half (+369 to +468) exhibited termination similar to wild-type. We tested Hrp1 prey interaction with *SNR82* bait RNA in the B3H assay, and while there was significantly more β-gal activity in the +insert versus -insert negative control (-Bait), the overall interaction was modest and not further explored in this context (**Fig. S4**). We conclude that both halves of the 200 bp *SNR82* terminator (+269 to +468) are necessary for full activity in yeast, but the upstream portion (+269 to +369) is more active than the downstream portion (+369 to +469) when partitioned.

**Figure 6.**
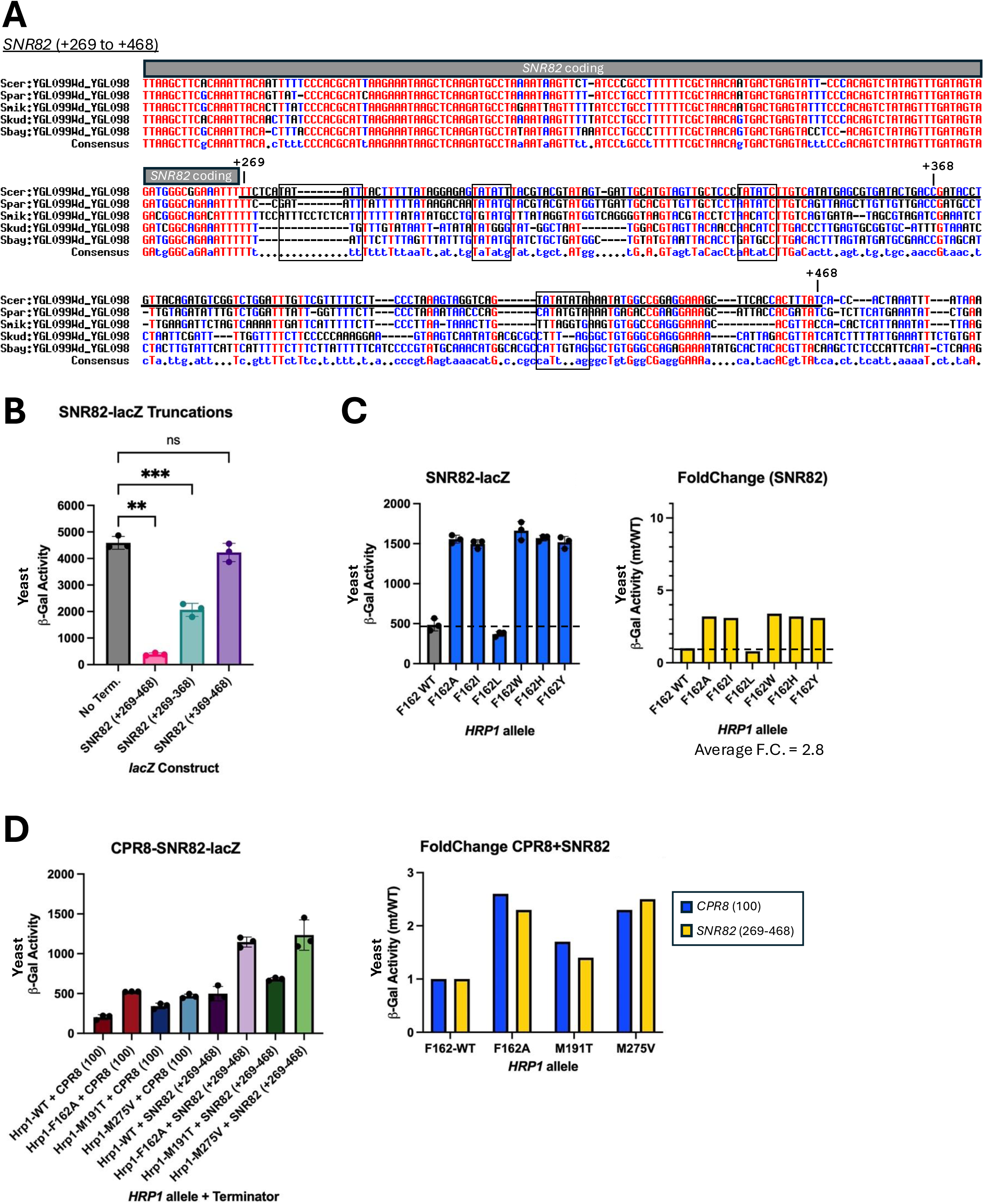
Characterizing Hrp1-dependent Pol II termination at the *SNR82* non-coding 3’-end terminator and the *CPR8* mRNA promoter-proximal terminator. **(A)** Yeast DNA downstream of the non-coding *SNR82* gene was aligned as described in Figure 1. Coding regions are indicated, and transcription terminator regions cloned into reporter genes are underlined, with numbering relative to the *SNR82* transcription start site (+1). **(B)** Yeast strains bearing full-length or truncated *SNR82* terminator-*lacZ* reporter gene plasmids were grown to saturation overnight at 30°C and diluted prior to cell lysis and incubation with ONPG substrate, using absorption at OD_600_ for cell density and OD_420_ for β-gal production. Experiments were performed in biological triplicate, and error bars show standard deviation. **(C)** Yeast strains bearing the *SNR82* (+269-468) terminator-*lacZ* reporter gene plasmid and HA-Hrp1 WT or F162 mutant plasmids were analyzed as described in panel (B). Fold-change was calculated by dividing mutant β-gal values by their respective wild-type β-gal values. **(D)** Yeast strains bearing the *CPR8*(100) or *SNR82*(+269-468) terminator-*lacZ* reporter gene plasmids and HA-Hrp1 WT, F162A, M191T, or M275V mutant plasmids were analyzed as described in panel (B).

### The *SNR82* and *CPR8* terminators are similarly sensitive to Hrp1 RRM mutants F162A, M191T, and M275V in yeast

Finally, we compared the sensitivity of *SNR82* 3’-end versus *CPR8* 5’-end terminators when the Hrp1 RRM is mutated in yeast cells. The pattern we observed for the *SNR82*-*lacZ* construct was different from the previously tested *GAL7* pA site and the *RAD3*, *MNR2*, and *CPR8* upstream terminators. For the *SNR82* terminator, the F162A, I, W, H, and Y mutants exhibited comparable levels of terminator readthrough, but surprisingly F162L was similar to wild-type levels (**Fig. 6C**). The *CPR8* and *SNR82* terminators were similarly sensitive to Hrp1 RRM mutants F162A, M191T, and M275V, exhibiting 2-3 fold higher levels of β-gal activity versus wild-type. We conclude that the differential impact of Hrp1 RRM mutants on the *SNR82* 3’-end terminator in yeast (Goguen and Brow 2023) versus the *CPR8* 5’-end terminator in bacteria (this study) is not due to inherent differences in RNA sensitivity.

## DISCUSSION

The importance of transcription attenuation in regulating gene expression has expanded from its original discovery in bacteria (Turnbough 2019), and most pertinent for this study, attenuation is a significant event in eukaryotic systems (Kamieniarz-Gdula and Proudfoot 2019; Rouvière et al. 2022; Bentley 2024; Grzechnik and Mischo 2025). Given the existence of several Pol II termination pathways, each of which are linked to RNA 3’-end formation, questions remain regarding the role of RNA-binding proteins in attenuator recognition. Here, we expanded our understanding of 5’-end attenuator RNA elements and their recognition by the Hrp1 protein across a variety of termination sites.

### 5’-end RNA attenuators in RAD3, MNR2, SNG1 yeast genes resemble 3’-end polyadenylation sites, including putative efficiency elements

In this study we identified readthrough mutations across several Pol II attenuators, many of which clustered in TA-rich DNA elements that become UA-rich RNA once transcribed. The 5’-end UA-rich attenuator elements strongly resemble pA site efficiency elements (EE), which are known binding sites for Hrp1 (consensus 5’-UAUAUA-3’) during mRNA 3’-end processing (Kessler et al. 1997; Valentini et al. 1999; Kaur et al. 2021). In many cases the TA-rich elements are phylogenetically conserved across different yeast species, suggesting an important biological function (Scannell et al. 2011). Hrp1 was sufficient for binding *RAD3* and *MNR2* attenuators in a B3H assay, and RNA mutations in UA-rich regions disrupted the interaction. Readthrough mutations in the *CPR8* terminator (T-94C, T-72A, A-71G) were insufficient to disrupt B3H interaction with Hrp1, but a mutant in a less conserved TA-rich element (T-103A) remains to be tested. In cases where RNA readthrough mutations in yeast translated to lower Hrp1-RNA interaction in the B3H assay, the impact was not a complete disruption as observed for the Hrp1-F162A mutant. This effect may be due to some remaining function at the mutated RNA site or Hrp1 binding to a neighboring non-mutated site. Similar to the *GAL7* 3’-end pA site, *RAD3* and *MNR2* 5’-end attenuator RNAs contain extended TA-rich elements. Hrp1 may bind cooperatively to longer EE elements based on a 2:1 Hrp1-RNA stoichiometry in chemical shift perturbation experiments as well as higher molecular weight complexes observed during elevated protein-RNA ratios (Perez-Canadillas 2006). The *GAL7*-mut(x5) allele created in this study contains five mutations spread across two tandem copies of the (UA)_6_ RNA element, which ablated the B3H interaction to near background levels. Similar multi-mutant approaches may be necessary to fully disrupt Hrp1 binding at attenuator regions.

In addition to Hrp1, it remains to be seen whether other RNA-binding proteins are involved in recognition of the *RAD3*, *MNR2*, and *SNG1* attenuators. The CFIA components Rna14 and Rna15 seem likely to be involved given their association with Hrp1/CFIB (Gross and Moore 2001; Leeper et al. 2010), and they have previously been shown to promote recognition of the *DEF1* “hybrid” attenuator (Whalen et al. 2018). Extended UA-rich elements have been implicated as supermotifs in NNS termination, including targets in snoRNAs (*SNR13*, *SNR33*) and cryptic unstable transcripts (*IMD2*, *URA2*) (Porrua et al. 2012). Notably for this study, there is an absence of attenuator readthrough mutations in regions matching the consensus binding sites for Nrd1 (5’-GUAA-3’, 5’-GUAG-3’) and Nab3 (5’-UCUU-3’) (Steinmetz and Brow 1998; Carroll et al. 2004; Creamer et al. 2011; Porrua et al. 2012). It is yet to be determined if the Hrp1-dependency of *RAD3*, *MNR2*, and *SNG1* attenuators is similar to the premature cleavage and polyadenylation (PCPA) observed in human mRNA introns (Vlasenok et al. 2023). Cryptic unstable transcripts have been documented in promoter-proximal regions for *MNR2* and *SNG1* (Neil et al. 2009), suggesting that attenuated transcripts from these genes are rapidly degraded through the TRAMP/exosome complex (Colin et al. 2011; Jensen et al. 2013). Hrp1 may coordinate Sen1 recruitment with limited influence for Nrd1/Nab3, as has been proposed for the *DEF1* attenuator (Whalen et al. 2018), or Hrp1 may interact directly with Pol II to influence termination (Wang et al. 2026). As a possible parallel to yeast Hrp1-dependent attenuation, the metazoan Restrictor complex prematurely terminates Pol II seemingly without transcript cleavage (Estell and West 2025), and its selectivity relies on sequence-specific recognition of RNA 5’-end motifs (Libri 2026; Polizzese et al. 2026).

### An important RNA-binding feature of the highly conserved Hrp1 RRM1 amino acid residue F162 relates to its aromatic property

The Hrp1 RRMs are highly conserved across a wide variety of RNA-binding proteins in yeast and other eukaryotes (Perez-Canadillas 2006; Goguen and Brow 2023), and our previous work identified substitutions in RRM1 F162 that were disruptive for attenuator recognition (Amodeo et al. 2022). Structural evidence of Hrp1 in complex with (UA)_4_ RNA suggests that the aromatic R-group of F162 stacks between the U5 and A6 bases, forming π-π interactions (Perez-Canadillas 2006; Barnwal et al. 2012). A genetic selection for Hrp1 readthrough mutations of the *SNR82* non-coding RNA terminator implicated F162 as part of a “3-bridge cluster” with F202 and F204, coordinated by a central M191 amino acid and expanding RNA contacts to include U7 (Goguen and Brow 2023). Our decision to test group one (F162A, F162I, F162L) and group two (F162W, F162H, F162Y) Hrp1 mutants was based on human alternative splicing factor Fox1, where residue F126 forms direct π-stacking interactions with bases U1 and G2 (Auweter et al. 2006). The Fox1-F126Y mutant behaved most similarly to the Fox1 wild-type protein, reducing affinity ∼2 fold when tested via *in vitro* binding assays, which is consistent with our observation that Hrp1-F162Y is the only viable yeast mutant in single copy. The group one F162A and F162I mutants were most detrimental to Fox1 function, reducing affinity by ∼1500-fold. The Fox1-F162L mutant was more intermediate in function with binding reduced ∼300-fold, suggesting that hydrophobic packing could substitute for an aromatic side chain if there was less steric hindrance. Interestingly, we observed near wild-type function for Hrp1-F162L when tested with the *SNR82* 3’-end RNA terminator but not 5’-end attenuator RNAs, suggesting a difference in RNA sequence context and/or protein binding partners. Additional studies will be necessary to understand the subtleties between Hrp1-dependent terminators, particularly at 5’-end protein-coding vs. 3’-end non-coding RNA targets.

### The bacterial 3-hybrid (B3H) system is a useful model to study RNA-binding proteins in the absence of other eukaryotic components

The bacterial 3-hybrid system was developed as an assay to confirm putative RNA-protein interactions *in vivo* and assess how mutations in RNA or protein affect the strength of the interaction (Stockert et al. 2022). Recent updates to the system have improved the isolation of target RNA from neighboring sequences as well as provided an expression system for RNAs lacking their own intrinsic terminators (Nguyen et al. 2025). In this study, we adopted the B3H system to study a eukaryotic protein-RNA interaction, optimizing RNA size, protein size, and growth conditions. In some cases, a robust and significant B3H interaction was not observed, including RNA targets from *SNG1*, *DEF1*, *CYC1*, and *SNR82*. Future optimization may include longer and/or different RNA segments, with RNA secondary structure prediction tested using RNAFold to optimize the accessibility of predicted binding sites (Gruber et al. 2008).

The B3H assay allowed us to show that Hrp1 is sufficient to bind attenuator RNAs in the absence of other yeast components as well as characterize critical Hrp1-RNA interfaces. B3H interactions for Hrp1 with target RNAs ranged from ∼1.5-fold to ∼3-fold above empty vector negative controls, which is similar to what has been observed for the bacterial protein Hfq with 5’-UTR regions (Nguyen et al. 2025). The Hrp1-dependency of *CPR8* 5’-end and *SNR82* 3’-end RNA terminators mostly concurred between yeast reporter and B3H assays, supporting direct Hrp1 contact with RNA. However, five Hrp1 mutants (M191T, I270T, D271G, M275V, and T280I) failed to disrupt the *CPR8* B3H interaction despite their observed terminator readthrough defect in yeast. It was previously suggested that M191T and M275V mutants alter “3-bridge cluster” formation, I270T and D271G change interdomain interactions, and T280I disrupts protein-protein interactions with other termination factors and/or the Pol II elongation complex (Goguen and Brow 2023). The discrepancy between some Hrp1 mutant responses in yeast versus bacterial systems may be an artifact of the B3H system, such as fusion of the prey protein with the RNAP ⍺-NTD. A more interesting explanation is that some Hrp1 mutants exhibit a gain-of-function, requiring yeast factors missing from the B3H system for the defect to become apparent. Interestingly, an Hrp1-R317G substitution in RRM2 was identified as a genetic suppressor of Pol II mutant Rpb3-K9E (Goguen et al. 2026). The Rpb3-K9E mutant causes premature termination of Pol II, and Hrp1-R317G may act as a gain-of-function mutant by stabilizing the Pol II elongation complex. In this way, the B3H system could be a useful tool to decipher protein-RNA versus protein-protein functions for eukaryotic RNA-binding factors.

## DATA AVAILABILITY

The authors affirm that all data necessary for confirming the conclusions of the article are present within the article, figures, and tables. Tables S1, S2, S3, and S4 contain detailed descriptions of all yeast strains, primers, and plasmids used in this study. Strains and plasmids are available upon request.

## ACKNOWLEDGEMENTS

We thank Katherine Berry, Pádraig Deighan, and Ann Hochschild for reagents, and Experimental Biology course students (Emmanuel College) for preliminary investigations.

---

**Table S1.**
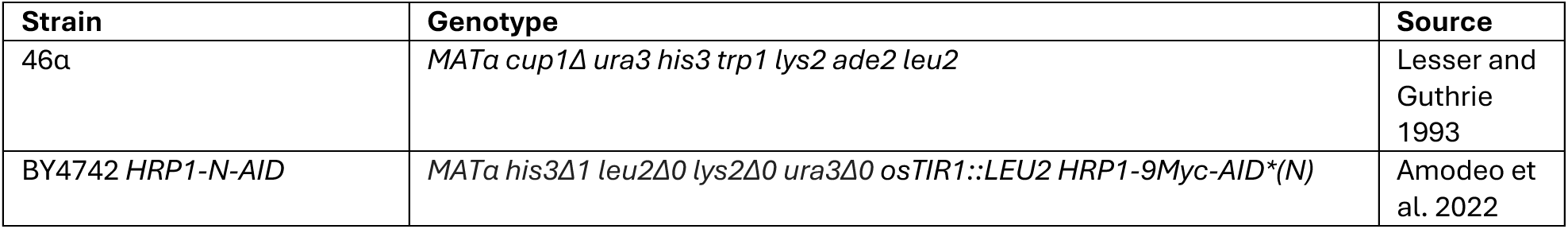
Yeast strains used in this study.

**Table S2.**
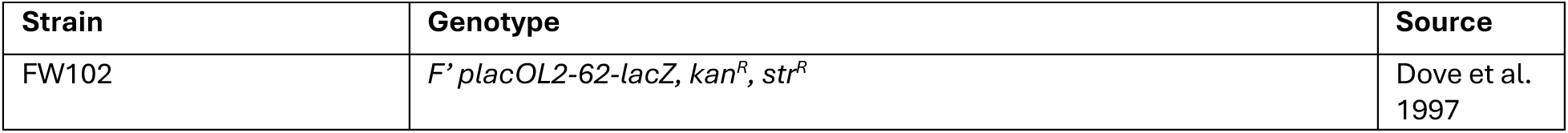
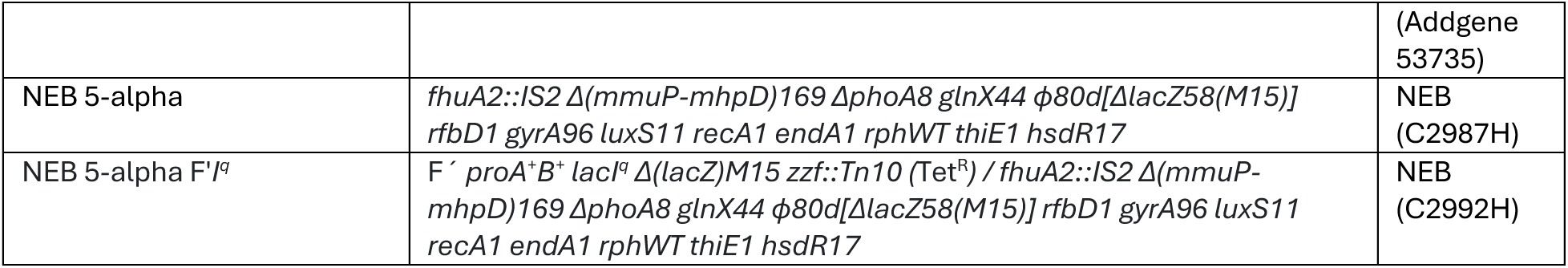
Bacterial strains used in this study.

**Table S3.**
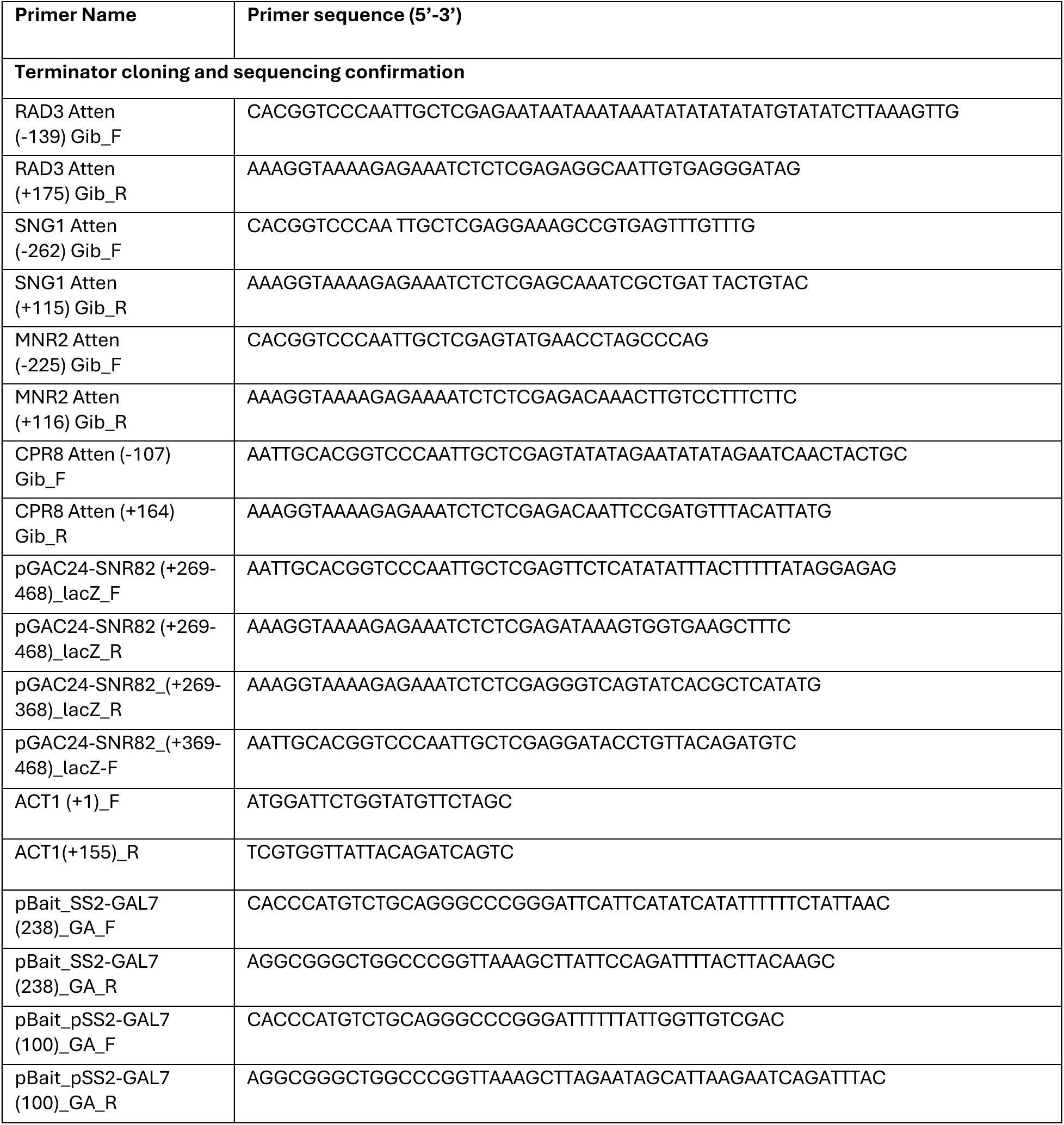
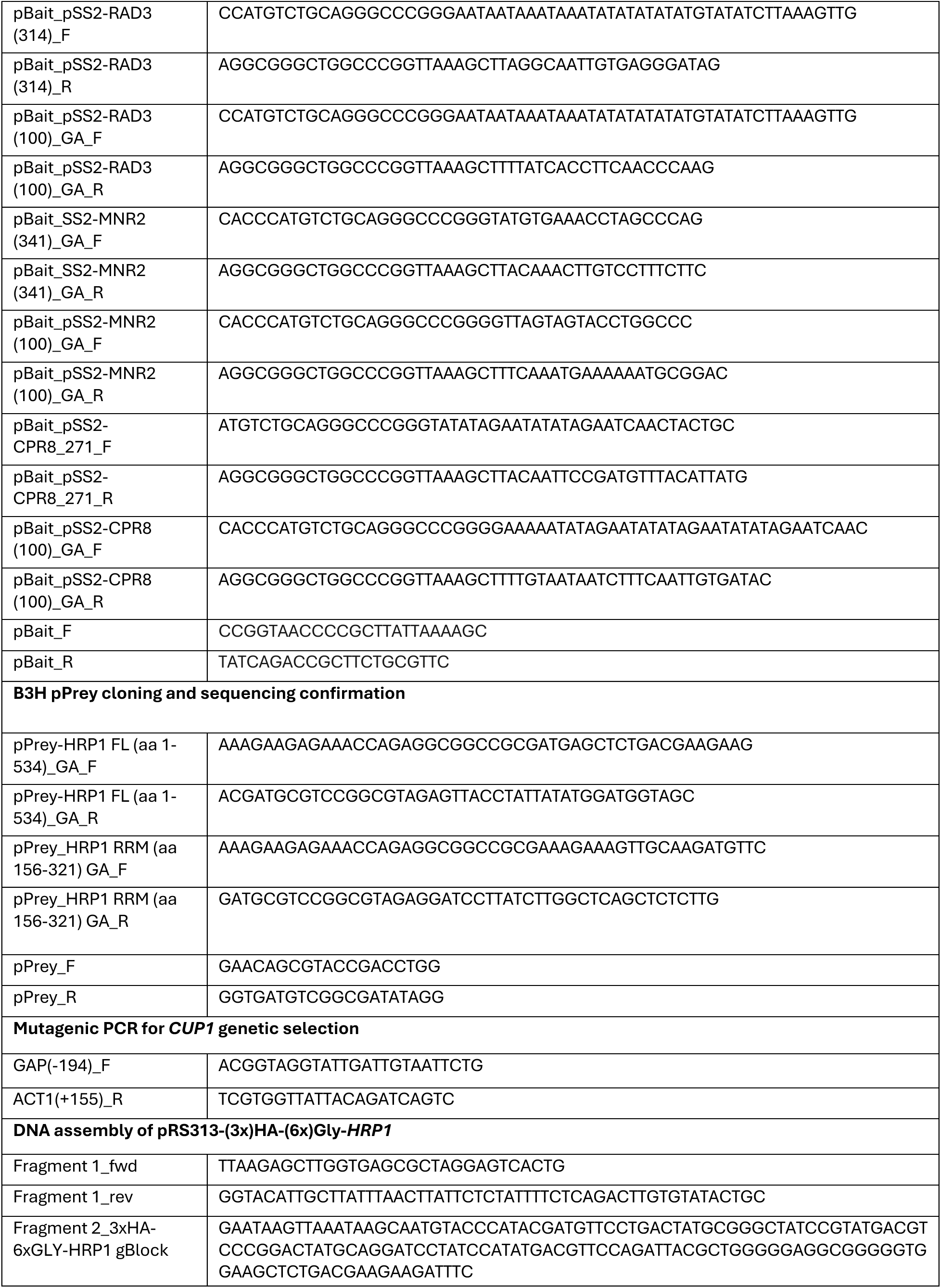
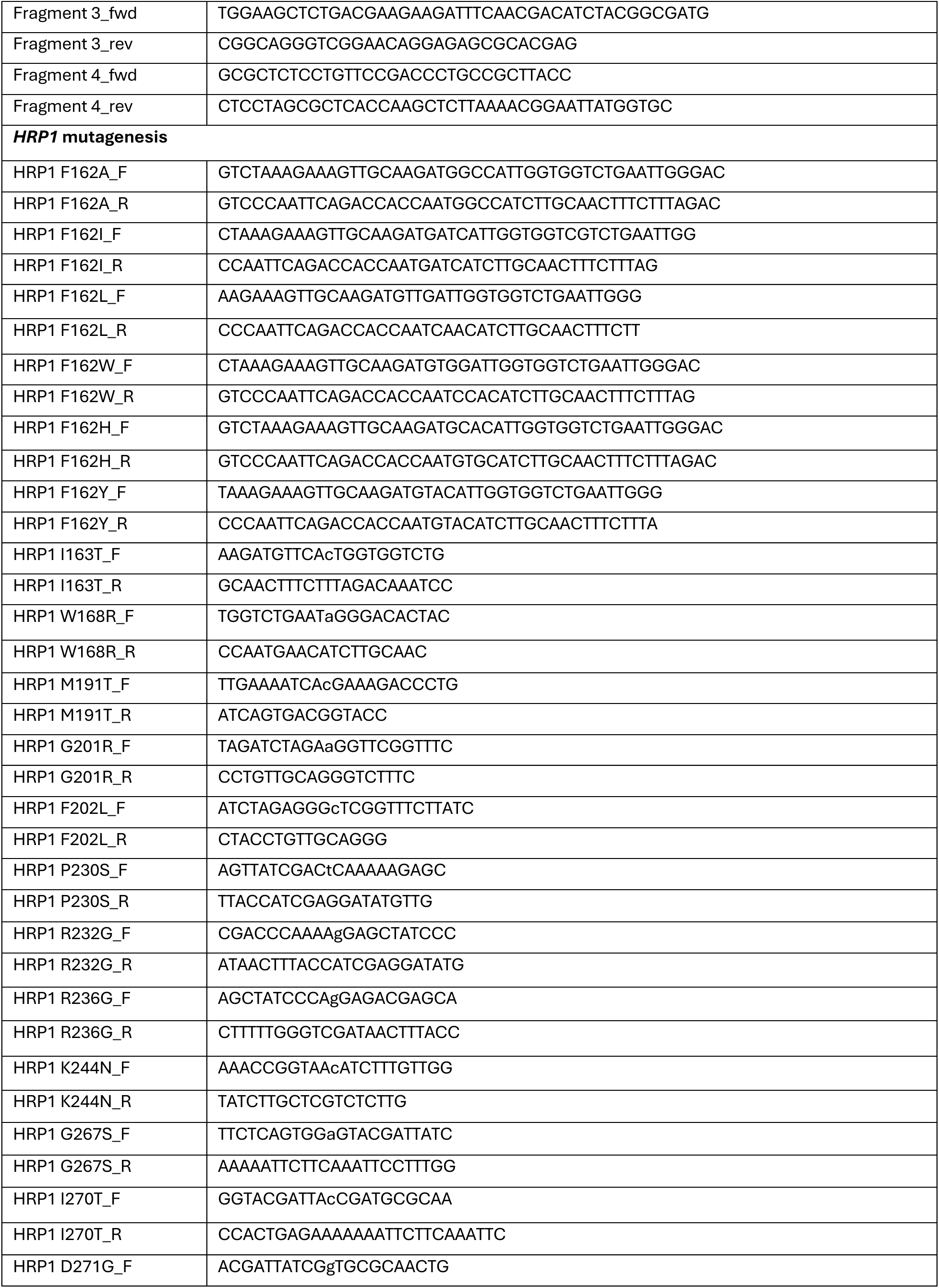
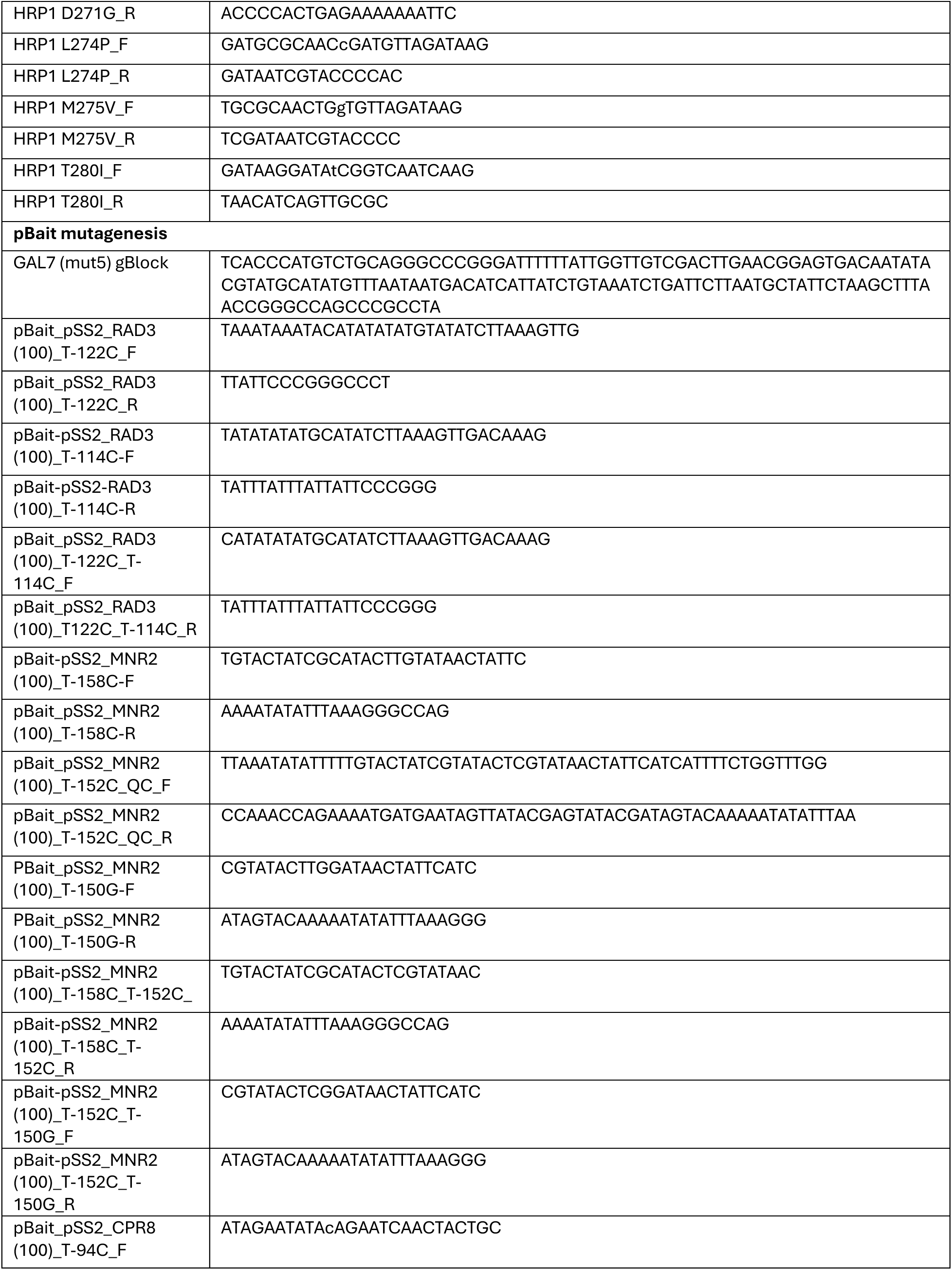
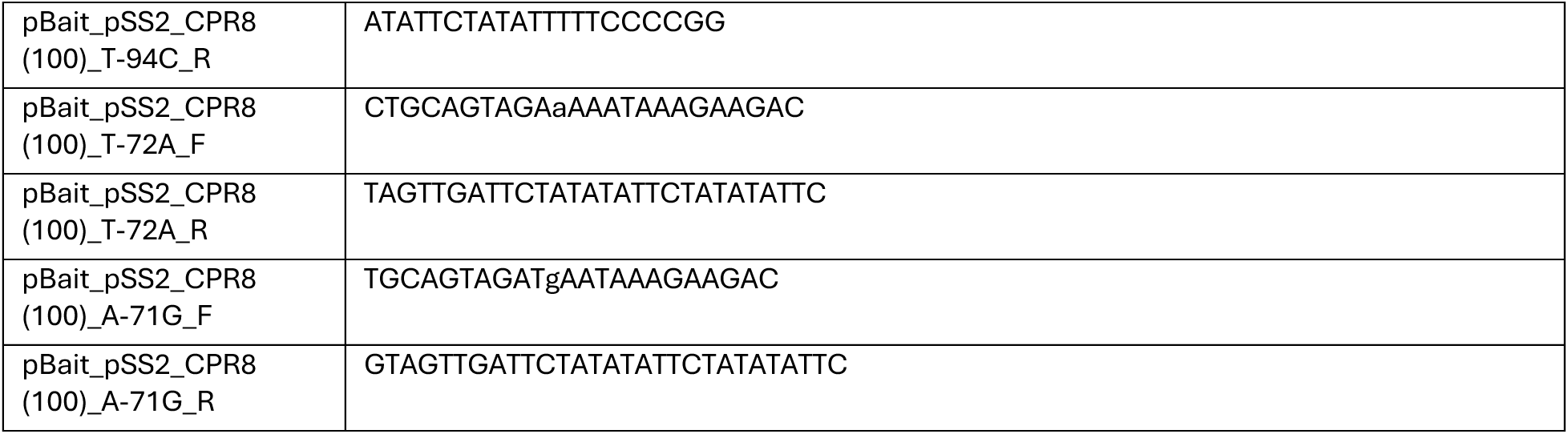
Primers used in this study.

**Table S4.**
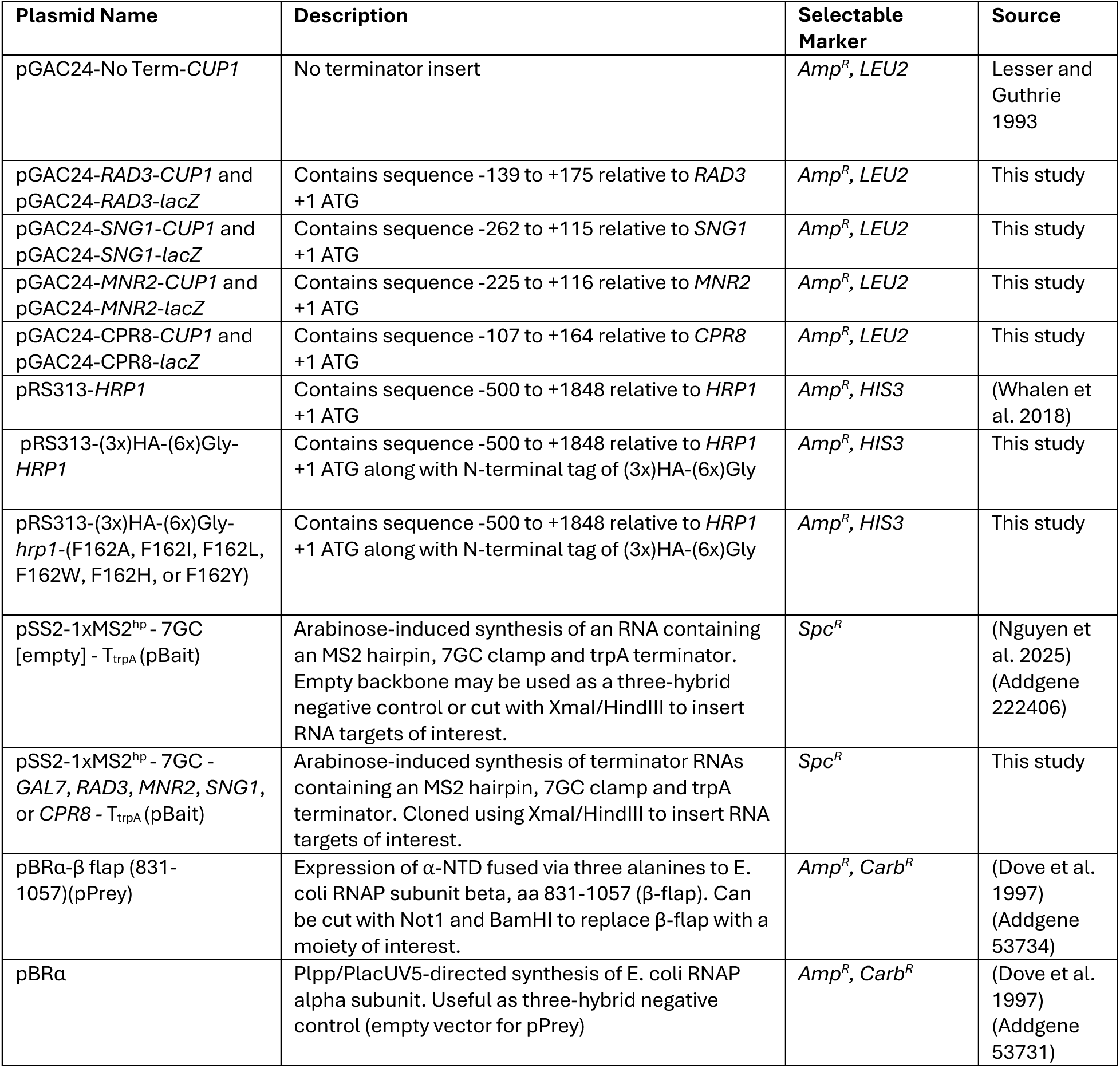
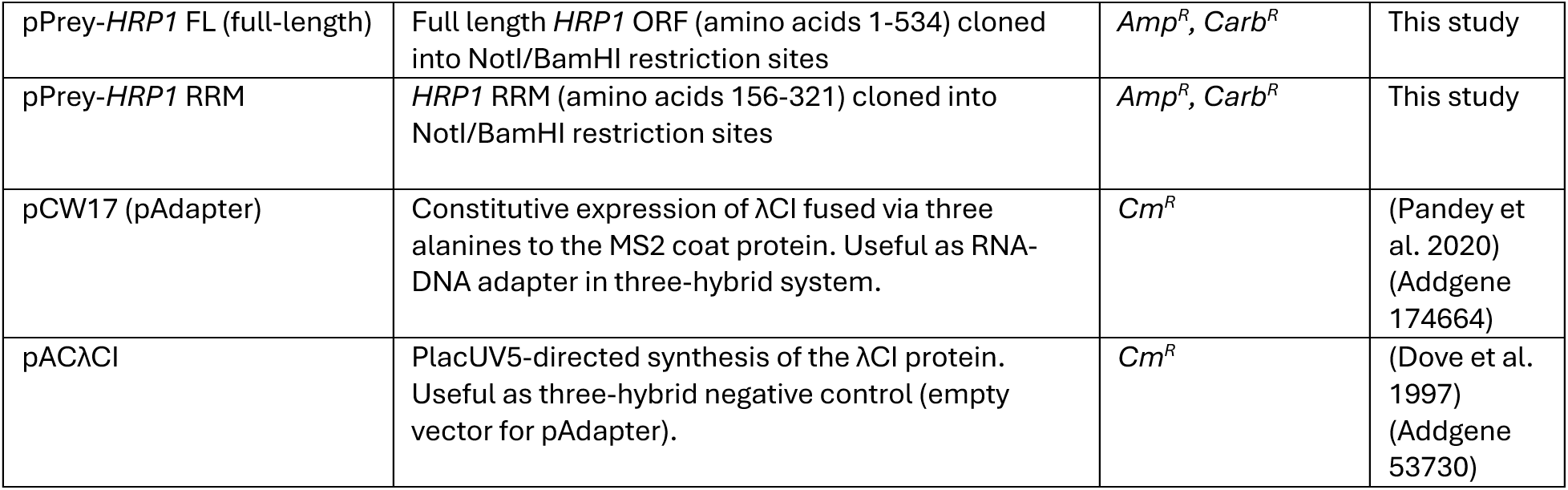
Plasmids used in this study.

| Plasmid Name | Description | Selectable Marker | Source |
| --- | --- | --- | --- |
| pGAC24-No Term-*CUP1* | No terminator insert | *Amp<sup>R</sup>*, *LEU2* | Lesser and Guthrie 1993 |
| pGAC24-*RAD3-CUP1* and pGAC24-*RAD3-lacZ* | Contains sequence -139 to +175 relative to *RAD3* +1 ATG | *Amp<sup>R</sup>*, *LEU2* | This study |
| pGAC24-*SNG1-CUP1* and pGAC24-*SNG1-lacZ* | Contains sequence -262 to +115 relative to *SNG1* +1 ATG | *Amp<sup>R</sup>*, *LEU2* | This study |
| pGAC24-*MNR2-CUP1* and pGAC24-*MNR2-lacZ* | Contains sequence -225 to +116 relative to *MNR2* +1 ATG | *Amp<sup>R</sup>*, *LEU2* | This study |
| pGAC24-*CPR8-CUP1* and pGAC24-*CPR8-lacZ* | Contains sequence -107 to +164 relative to *CPR8* +1 ATG | *Amp<sup>R</sup>*, *LEU2* | This study |
| pRS313-*HRP1* | Contains sequence -500 to +1848 relative to *HRP1* +1 ATG | *Amp<sup>R</sup>*, *HIS3* | (Whalen et al. 2018) |
| pRS313-(3x)HA-(6x)Gly-*HRP1* | Contains sequence -500 to +1848 relative to *HRP1* +1 ATG along with N-terminal tag of (3x)HA-(6x)Gly | *Amp<sup>R</sup>*, *HIS3* | This study |
| pRS313-(3x)HA-(6x)Gly-*hrp1*-(F162A, F162I, F162L, F162W, F162H, or F162Y) | Contains sequence -500 to +1848 relative to *HRP1* +1 ATG along with N-terminal tag of (3x)HA-(6x)Gly | *Amp<sup>R</sup>*, *HIS3* | This study |
| pSS2-1xMS2<sup>hp</sup> - 7GC [empty] - *T<sub>trpA</sub>* (pBait) | Arabinose-induced synthesis of an RNA containing an MS2 hairpin, 7GC clamp and *trpA* terminator. Empty backbone may be used as a three-hybrid negative control or cut with *XmaI*/*HindIII* to insert RNA targets of interest. | *Spc<sup>R</sup>* | (Nguyen et al. 2025) (Addgene 222406) |
| pSS2-1xMS2<sup>hp</sup> - 7GC - *GAL7*, *RAD3*, *MNR2*, *SNG1*, or *CPR8* - *T<sub>trpA</sub>* (pBait) | Arabinose-induced synthesis of terminator RNAs containing an MS2 hairpin, 7GC clamp and *trpA* terminator. Cloned using *XmaI*/*HindIII* to insert RNA targets of interest. | *Spc<sup>R</sup>* | This study |
| pBRα-β flap (831-1057)(pPrey) | Expression of α-NTD fused via three alanines to E. coli RNAP subunit beta, aa 831-1057 (β-flap). Can be cut with *NotI* and *BamHI* to replace β-flap with a moiety of interest. | *Amp<sup>R</sup>*, *Carb<sup>R</sup>* | (Dove et al. 1997) (Addgene 53734) |
| pBRα | Plpp/PlacUV5-directed synthesis of E. coli RNAP alpha subunit. Useful as three-hybrid negative control (empty vector for pPrey) | *Amp<sup>R</sup>*, *Carb<sup>R</sup>* | (Dove et al. 1997) (Addgene 53731) |
| pPrey-*HRP1* FL (full-length) | Full length *HRP1* ORF (amino acids 1-534) cloned into NotI/BamHI restriction sites | *Amp<sup>R</sup>*, *Carb<sup>R</sup>* | This study |
| pPrey-*HRP1* RRM | *HRP1* RRM (amino acids 156-321) cloned into NotI/BamHI restriction sites | *Amp<sup>R</sup>*, *Carb<sup>R</sup>* | This study |
| pCW17 (pAdapter) | Constitutive expression of λCI fused via three alanines to the MS2 coat protein. Useful as RNA-DNA adapter in three-hybrid system. | *Cm<sup>R</sup>* | (Pandey et al. 2020) (Addgene 174664) |
| pACλCI | PlacUV5-directed synthesis of the λCI protein. Useful as three-hybrid negative control (empty vector for pAdapter). | *Cm<sup>R</sup>* | (Dove et al. 1997) (Addgene 53730) |

## Supplementary Figures

**Figure S1.**
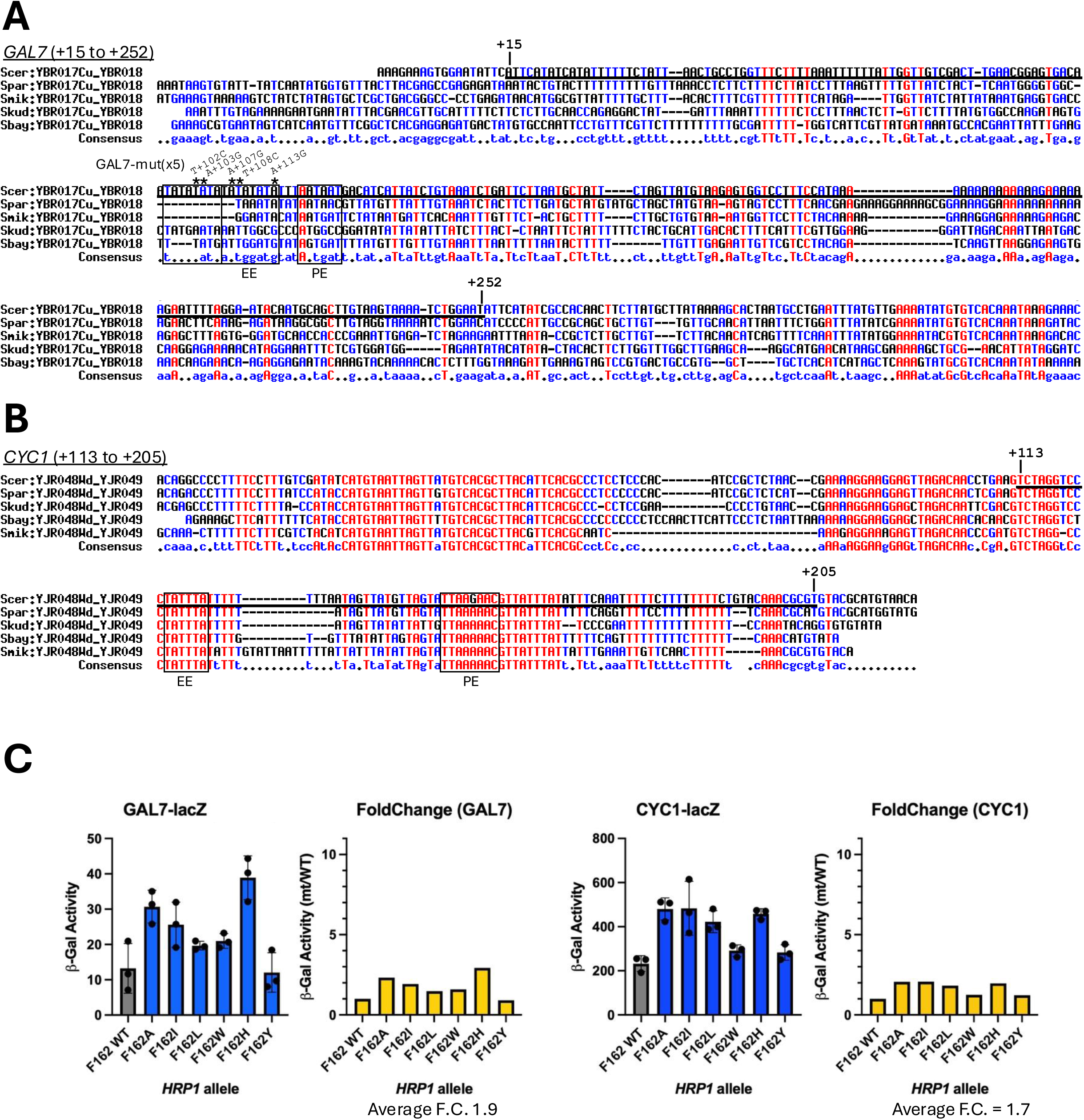
DNA sequence conservation and Hrp1-dependence of *GAL7* and *CYC1* 3’-end pA site terminators. Yeast DNA downstream of the protein-coding genes **(A)** *GAL7* and **(B)** *CYC1* was aligned as described in Figure 2. The base numbering is relative to the stop codon. Regions cloned into *lacZ* reporter genes are underlined. Consensus efficiency elements (EE) and positions elements (PE) are boxed. **(C)** Yeast strains bearing HA-Hrp1 WT and mutant plasmids and terminator-*lacZ* reporter gene plasmids were grown to saturation overnight at 30°C and diluted prior to cell lysis and incubation with ONPG substrate, using absorption at OD_600_ for cell density and OD_420_ for β-gal production. Experiments were performed in biological triplicate, and error bars show standard deviation. Fold-change was calculated by dividing mutant β-gal values by their respective wild-type β-gal values.

**Figure S2.**
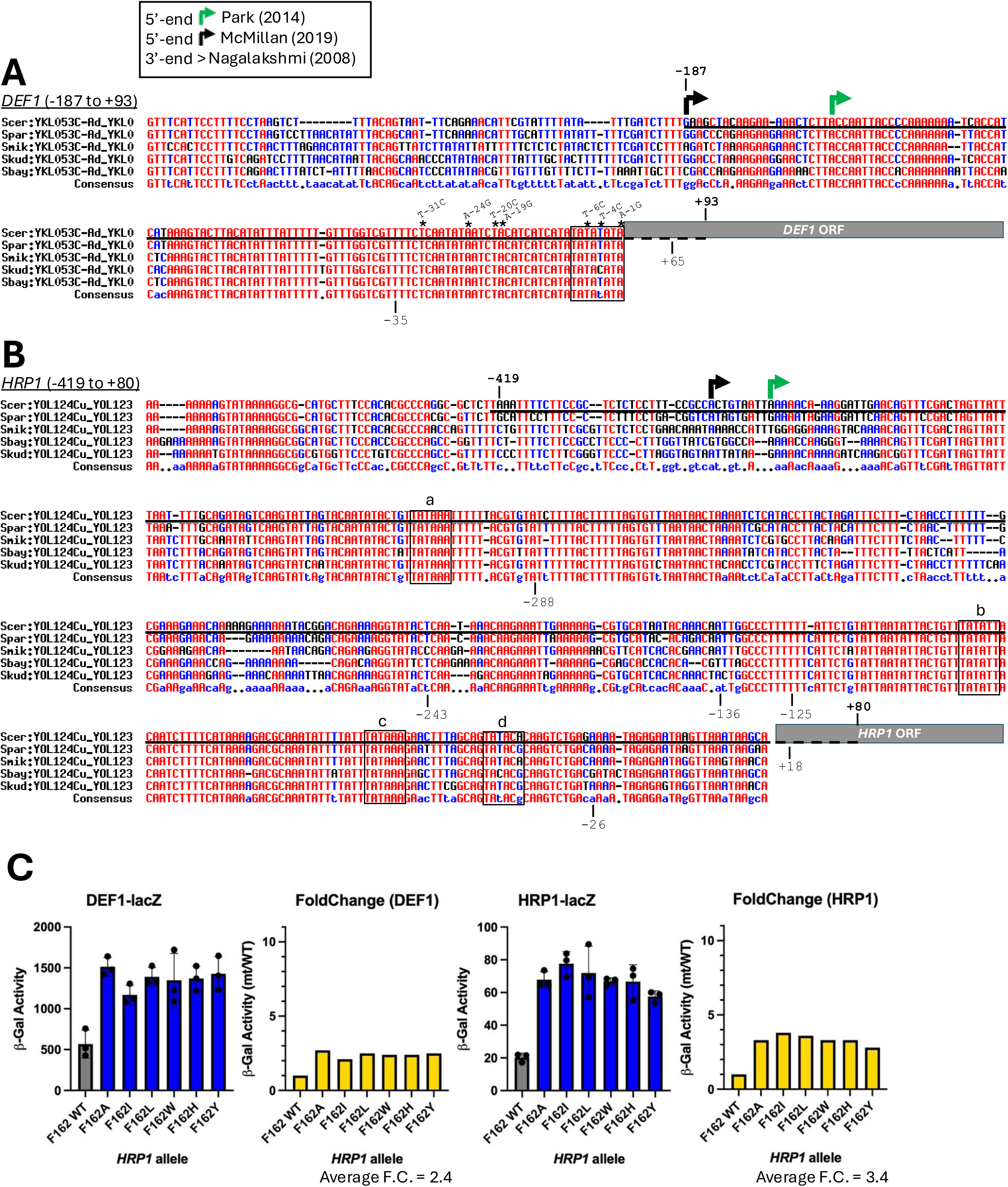
DNA sequence conservation and Hrp1-dependence of *DEF1* and *HRP1* attenuators. **(A)** and **(B)** Yeast DNA upstream of the protein-coding genes *DEF1* and *HRP1* was aligned as described in Figure 2. The 5’-end transcription start sites are indicated with arrows (black and green), and transcription terminator regions cloned into reporter genes are underlined, with numbering relative to the +1 <u>A</u>TG start codon. Putative TA-rich efficiency elements are boxed. Representative copper-resistant point mutants for *DEF1* are indicated with asterisks and numbered relative to the respective gene +1 <u>A</u>TG start codon (Whalen et al. 2018). **(C)** Yeast strains bearing HA-Hrp1 WT and mutant plasmids and terminator-*lacZ* reporter gene plasmids were grown to saturation overnight at 30°C and diluted prior to cell lysis and incubation with ONPG substrate, using absorption at OD_600_ for cell density and OD_420_ for β-gal production. Experiments were performed in biological triplicate, and error bars show standard deviation. Fold-change was calculated by dividing mutant β-gal values by their respective wild-type β-gal values.

**Figure S3.**
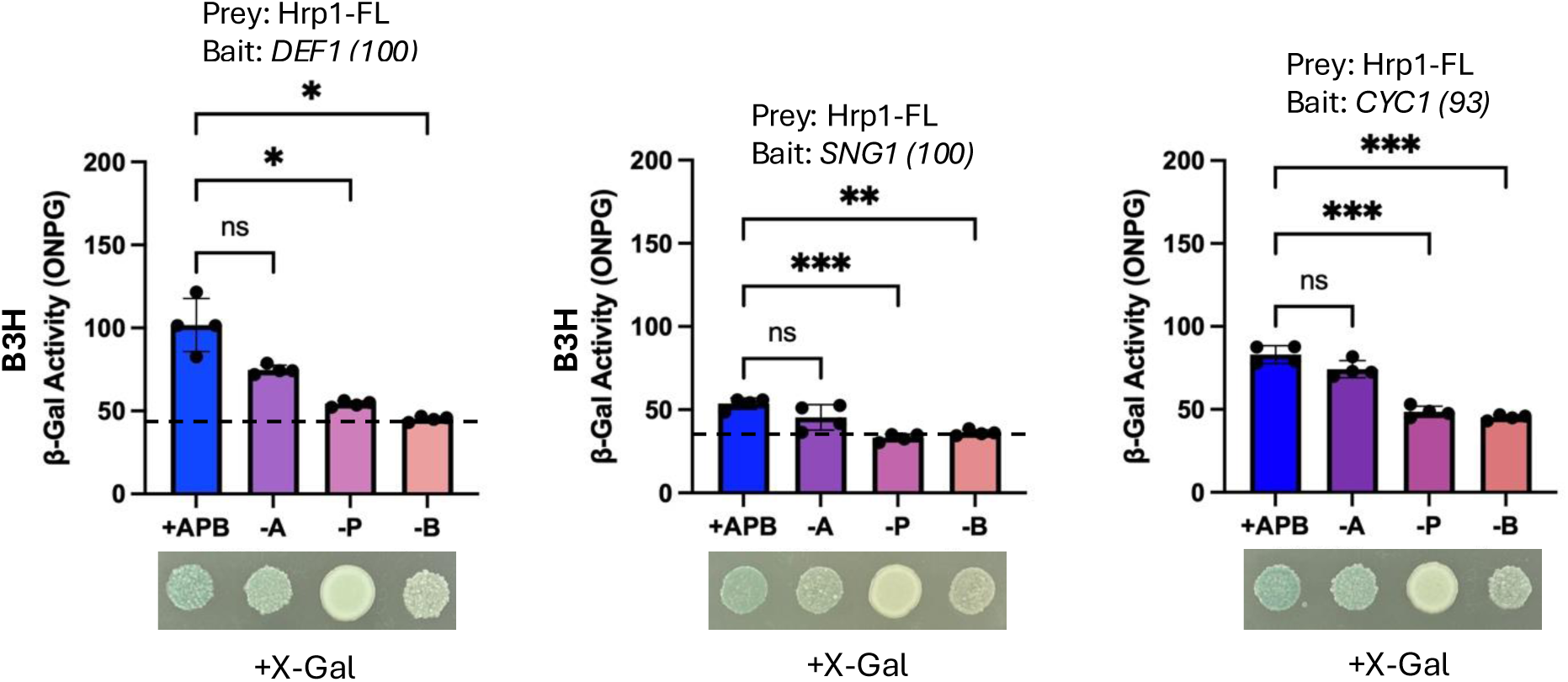
Bacterial 3-hybrid (B3H) assay fails to reliably detect Hrp1 protein interaction with 5’-end attenuators (*DEF1*, *SNG1*) and the *CYC1* 3’-end pA site. Bacterial strains bearing the indicated B3H reporter gene plasmids were grown to saturation overnight at 37°C, diluted back and grown to exponential phase, followed by cell lysis and incubation with ONPG substrate, using absorption at OD_600_ for cell density and OD_420_ for β-gal production. Experiments were performed using 3-4 biological replicates, and error bars show standard deviation. +APB = +Adapter, +Prey, +Bait. -A = -Adapter. -P = - Prey. -B = -Bait. Asterisks indicate statistical significance by Welch’s ANOVA (*p<0.05, **p<0.01, ***p<0.001, ns – not significant, p≥0.05). The same bacterial strains were diluted and spotted onto selective media containing X-gal substrate.

**Figure S4.**
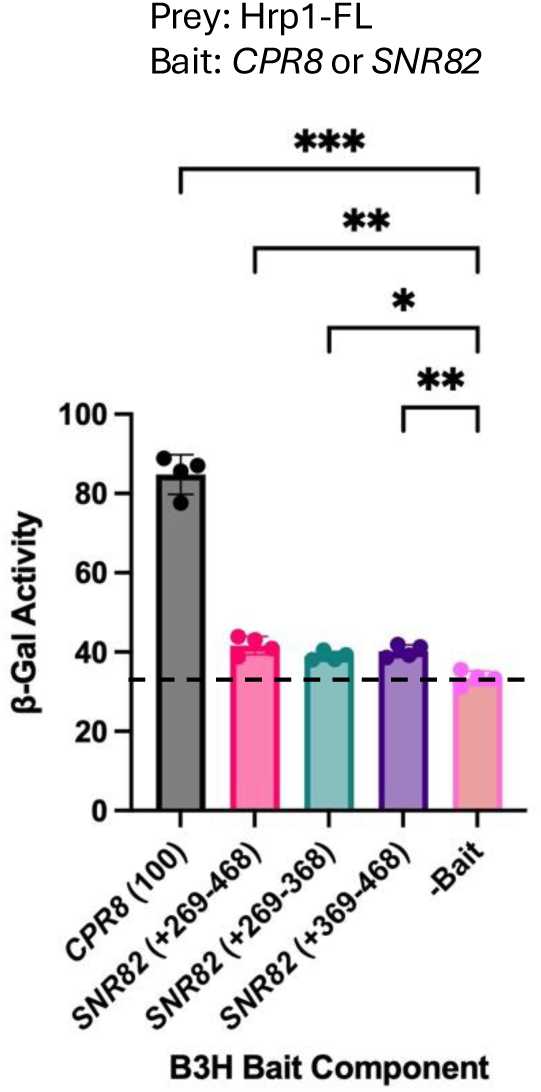
Bacterial 3-hybrid (B3H) assay detects a modest Hrp1 protein interaction with the *SNR82* 3’-end RNA terminator. Bacterial strains bearing the indicated B3H reporter gene plasmids were grown to saturation overnight at 37°C, diluted back and grown to exponential phase, followed by cell lysis and incubation with ONPG substrate, using absorption at OD_600_ for cell density and OD_420_ for β-gal production. -Bait is lacking RNA<u>X</u> and serves as a negative control. The *CPR8* (100) RNA bait serves as a positive control. Experiments were performed using 3-4 biological replicates, and error bars show standard deviation. Asterisks indicate statistical significance by Welch’s ANOVA (*p<0.05, **p<0.01, ***p<0.001, ns – not significant, p≥0.05).

